# Faster processing of moving compared to flashed bars in awake macaque V1 provides a neural correlate of the flash lag illusion

**DOI:** 10.1101/031146

**Authors:** Manivannan Subramaniyan, Alexander S. Ecker, Saumil S. Patel, R. James Cotton, Matthias Bethge, Xaq Pitkow, Philipp Berens, Andreas S. Tolias

## Abstract

When the brain has determined the position of a moving object, due to anatomical and processing delays, the object will have already moved to a new location. Given the statistical regularities present in natural motion, the brain may have acquired compensatory mechanisms to minimize the mismatch between the perceived and the real position of a moving object. A well-known visual illusion — the flash lag effect — points towards such a possibility. Although many psychophysical models have been suggested to explain this illusion, their predictions have not been tested at the neural level, particularly in a species of animal known to perceive the illusion. Towards this, we recorded neural responses to flashed and moving bars from primary visual cortex (V1) of awake, fixating macaque monkeys. We found that the response latency to moving bars of varying speed, motion direction and luminance was shorter than that to flashes, in a manner that is consistent with psychophysical results. At the level of V1, our results support the differential latency model positing that flashed and moving bars have different latencies. As we found a neural correlate of the illusion in passively fixating monkeys, our results also suggest that judging the instantaneous position of the moving bar at the time of flash — as required by the postdiction/motion-biasing model — may not be necessary for observing a neural correlate of the illusion. Our results also suggest that the brain may have evolved mechanisms to process moving stimuli faster and closer to real time compared with briefly appearing stationary stimuli.

**New and Noteworthy:** We report several observations in awake macaque V1 that provide support for the differential latency model of the flash lag illusion. We find that the equal latency of flash and moving stimuli as assumed by motion integration/postdiction models does not hold in V1. We show that in macaque V1, motion processing latency depends on stimulus luminance, speed and motion direction in a manner consistent with several psychophysical properties of the flash lag illusion.

## Introduction

Moving objects in nature typically follow smooth, predictable trajectories, potentially enabling the brain to minimize or compensate for motion processing delays. The flash lag illusion has kindled much interest among neuroscientists as it is thought to provide a window into the neural mechanisms of localizing moving objects. In this illusion, observers report that a moving bar is located ahead of an aligned flash (Fig. 1A) (Mackay 1958; Nijhawan 1994). While this phenomenon has been studied extensively at the behavioral level, its underlying neural mechanisms are poorly understood.

**Fig. 1.**
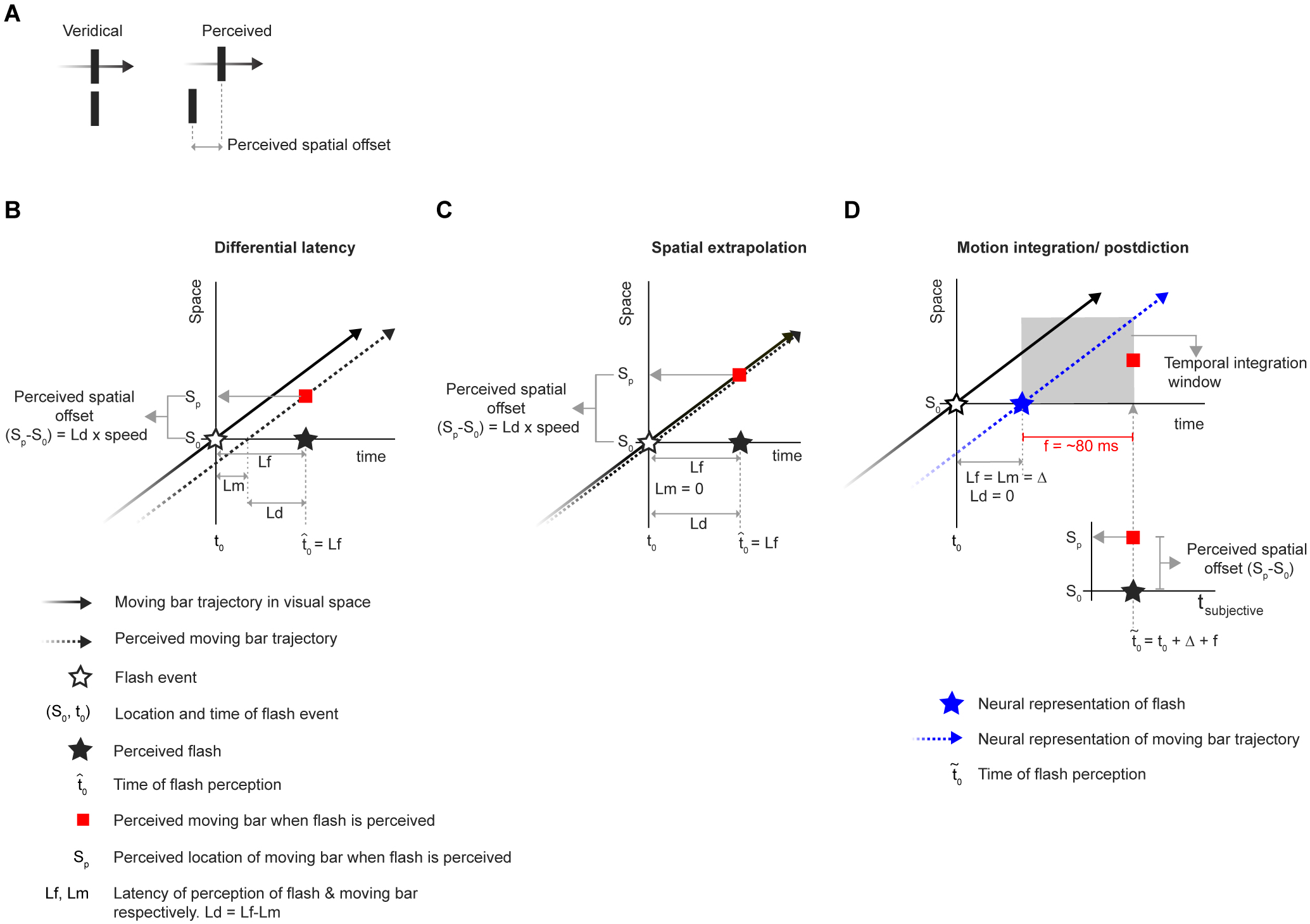
Models of flash lag illusion. ***A***, When a flash (bottom) is presented aligned to a moving bar (top), observers perceive the moving bar to be located further along the motion trajectory at the moment when they perceive the flash. Left: veridical locations; right: perceived locations. Since the flash appears to spatially lag behind the moving bar, this phenomenon has been called the flash lag illusion. In all panels *(**A-D**)*, motion is assumed to have started long before the flash event. ***B-D***, Illustration of flash lag effect as explained by different models. ***B***, Differential latency model. The flash (open star) presented at time tc at location S0 is perceived with a latency of Lf. Since the model assumes a shorter motion latency (Lm < Lf), when the flash is perceived (filled star), the moving bar located (at Sp, red square) further along the trajectory is perceived to be aligned with flash. ***C***, In the spatial extrapolation model, the moving bar is perceived ahead of the flash due to the longer latency of flash like in ***B*** except that the motion latency is assumed to be zero. ***D***, Illustration of the postdiction model, adapted from Fig. 2 of Rao et al (2001). In this model, the nervous system completes processing the flash at t_0_+Δ at which point the motion integration that lasts for a duration of *f* is triggered. Although in the other models the flash is perceived at t_0_+Δ, postdiction model claims that the perception of the flash is delayed until t_0_+Δ+f at which point the motion integration based moving bar position estimation is completed (Rao et al. 2001). Hence, although in the external time, motion integration based moving bar position is obtained at time t_0_+Δ+f and flash representation is completed earlier at t_0_+Δ, in the subjective time, they are perceived simultaneous, giving raise to the perceived spatial offset.

In an initial attempt to explain this illusion, it was posited that the brain extrapolates the position of moving stimuli to an extent that compensates its own processing delays (Nijhawan 1994). Since then, many alternative models have been proposed. These diverse models (Fig. 1B-D), reviewed extensively elsewhere (Eagleman and Sejnowski 2007; Öğmen et al. 2004), have pointed towards equally diverse neural mechanisms which range from simple bottom-up explanations such as latency differences to high level top-down mechanisms such as attention and feedback (Bachmann and Poder 2001; Baldo and Klein 1995; Brenner and Smeets 2000; Eagleman and Sejnowski 2007; Krekelberg and Lappe 2000; Patel 2000; Purushothaman et al. 1998; Sheth et al. 2000; Whitney and Murakami 1998). For example, the differential latency model (Purushothaman et al. 1998; Whitney and Murakami 1998) maintains that moving stimuli are processed faster compared to flashed ones, leading to the perception of flashes temporally coinciding with a moving bar further along its trajectory. Alternatively, the motion-biasing model (Eagleman and Sejnowski 2007; Rao et al. 2001) argues that motion signals after the detection of flash event affect position representation and judgments, such that observers report a misalignment between a flash and a moving stimulus. There has also been a recent attempt to subsume all these models into a single theoretical framework treating the flash lag effect as a probabilistic motion-based predictive shift (Khoei et al. 2017).

Most models of the flash lag illusion are formulated at the psychophysical level. At this level of abstraction, the three most prominent models (Fig. 1B-D) of the illusion differ in their prediction for the relative latencies of flashed and moving stimuli (Fig. 2). In this context, the latency or “representation delay” refers to the time interval between stimulus appearance at a particular location in the physical world and the emergence of neural activity corresponding to the reported perception of the stimulus location. The spatial extrapolation model, as it posits full compensation for neural delays (Nijhawan 1994), would predict zero representation delay for motion; the differential latency model predicts shorter latency for motion (Patel 2000; Purushothaman et al. 1998; Whitney and Murakami 1998) and the postdiction model assumes equal latency for flash and moving stimuli (Eagleman and Sejnowski 2007; Rao et al. 2001). The models do not specify in which part of the brain one would observe such predicted latency differences of stimulus representations. Here, to systematically investigate the neural mechanisms of the flash lag illusion and to test predictions of the psychophysical models, we measured the latencies or representation delays of flashed and moving stimuli in primary visual cortex (V1) of awake, fixating macaques. This allows us to determine the contribution of early visual processing towards the illusion. Note that, at the level of V1, the term latency or “representation delay” refers to the time interval between stimulus appearance at a particular location in the physical world and the time of stimulus-evoked activity in V1 at which a decoder or a downstream processing region can obtain the best estimate of the stimulus location (see Methods).

**Fig. 2.**
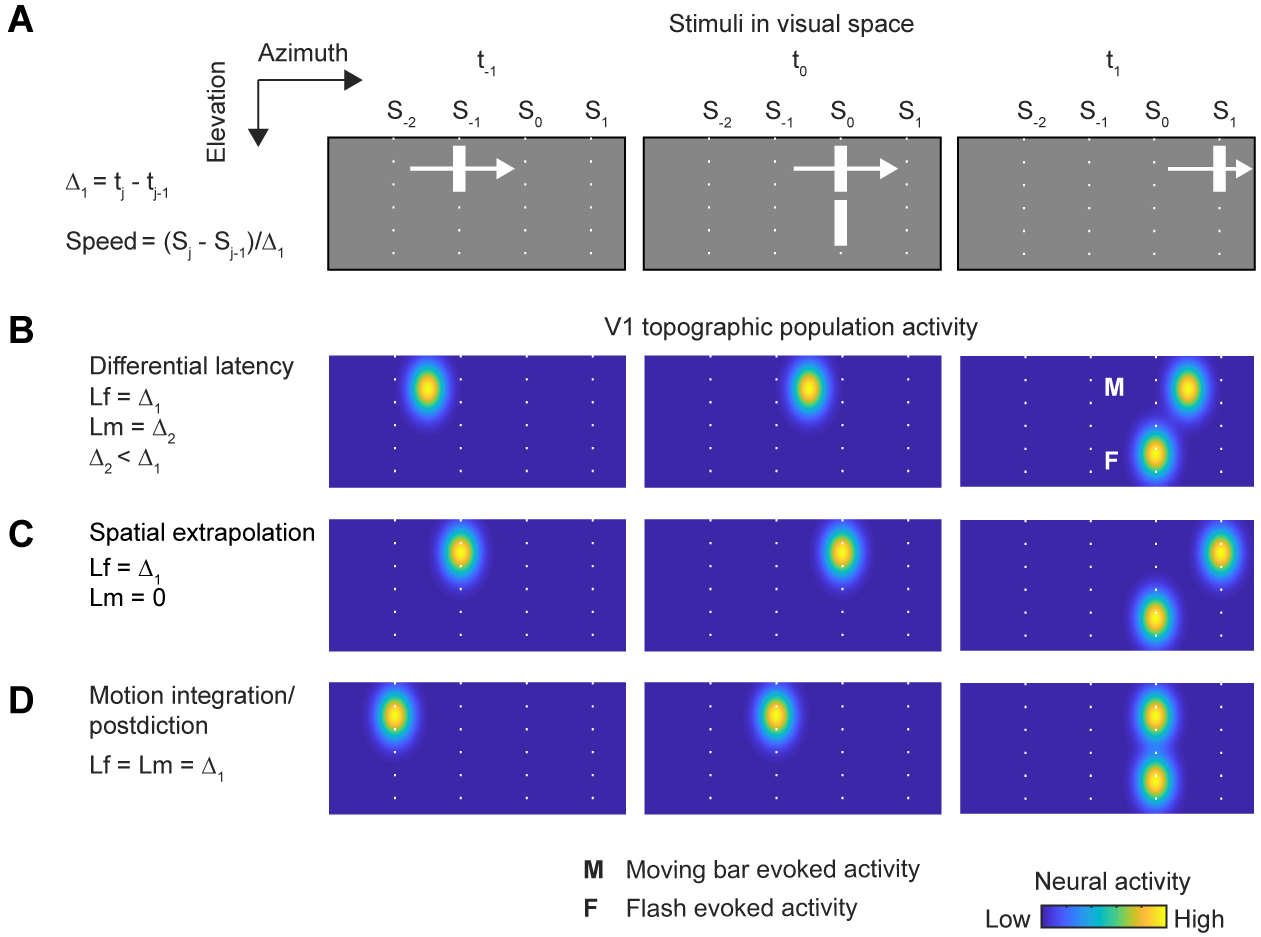
Predictions of models of flash lag illusion for V1 population activity for continuous motion condition with identical flash and moving bar luminance. In all panels *(**A-D**)*, motion is assumed to have started long before the flash event. ***A***, Illustration of hypothetical visual stimuli that generate the population activity in V1 as predicted by different models (***B-D***). For simplicity, the stimulus positions are shown at just three time instances t_−1_, t_0_ and t_1_. The time the moving bar takes (Δ_1_ = t_j_ − t_j−1_ ms) to traverse from S_j_ to S_j−1_ is also set to be equal to the latency of population activity peak for flash *(**B-D**). **B-D***, Illustrations showing predicted topographically organized V1 population neural response to stimuli depicted in matching panels in ***A***. The flash is assumed to be fully represented in V1 when the population hill reaches its peak activity at S_0_. In the moving bar condition, a fully developed population activity hill (white label ‘**M**’) representing some position, continuously translates following the motion trajectory. Hence which position of moving bar caused an activity hill at a given instant will depend on the motion population response peak latency. For all models, the neural representation of flash (white label ‘**F**’) in V1 is delayed by the same duration Δ_1_. The models differ in the neural representation delays of moving bar as seen at time instant t1: the differential latency model (***B***) predicts that the population hill will spatially lag behind the moving bar but will be shifted along the motion direction relative to flash population hill. The spatial extrapolation model (***C***) predicts a similar shift of the motion population hill relative to flash. However, the motion population hill does not spatially lag behind the moving bar. The postdiction model (***D***) assumes identical latency for flash and moving bar neural representations - hence the population hills will be aligned.

The few physiological studies that have explored the neural mechanisms of the illusion found a shorter latency for motion signals compared to flashes in the rabbit and salamander retina (Berry et al. 1999), cat LGN (Orban et al. 1985) and cat V1 (Jancke et al. 2004b), providing evidence for a bottom up latency difference between flashes and moving stimuli. Although these studies provide valuable hints at plausible neural mechanisms of the flash lag illusion, they were done either *in vitro* or in anesthetized animals and it is unknown if the animals used in these studies actually perceive the illusion.

We previously showed that, similar to humans, macaque monkeys perceive the flash lag illusion (Subramaniyan et al. 2013). Hence, we performed the physiological experiments in awake macaque monkeys, which allowed us to directly test the predictions of different models of the illusion at the level of V1 neural representation of flash and moving stimuli (Fig. 2). Specifically, we estimated the latency of the stimulus representations by two different methods, one based on multiunit response peak times and the other based on probabilistic decoding of simultaneously recorded single- and multiunit population activity. Crucially, we measured the dependence of latency on different stimulus parameters — speed, luminance and direction of motion — to test if the resulting changes in neural responses accounted for the corresponding changes in perception. Under all these manipulations, neural latency differences between flash and motion in V1 explained a large part of the psychophysically measured perceived spatial offsets. Thus, our results show that even at the very first cortical visual information processing stage a neural correlate of the illusion can be observed, providing mechanistic constraints on the models of the flash lag illusion.

## Materials and Methods

### Subjects

Four male macaque (*Macaca mulatta*) monkeys (A, CH, CL and L) weighing 8, 9, 12 and 9.5 kg respectively and aged 10, 8, 8 and 8 years respectively, were used in the physiological experiments. Cranial head post and scleral search coil were implanted in each monkey using standard aseptic surgical procedures. All animal procedures were approved by the Institutional Animal Care and Use Committee of Baylor College of Medicine and followed NIH regulations. Two of the authors (MS and SP) participated in psychophysical experiments following procedures approved by the Institutional Review Board of Baylor College of Medicine.

### Electrophysiological recording and data processing

We used chronically implanted tetrode arrays for recording neural activity from monkeys A, CL and CH as described previously (Ecker et al. 2010; Tolias et al. 2007). Briefly, in each monkey, we implanted chronically, arrays of 24 tetrodes on the left hemisphere over the operculum in area V1. The tetrodes were custom built from Nichrome or Platinum/Iridium wires. We implanted a 96-electrode microelectrode array (‘Utah’ array, Blackrock Microsystems, Salt Lake City, UT, USA) over area V1 on the right hemisphere in monkey L. For both tetrode arrays and Utah array, the neural signals were pre-amplified at the head-stage by unity gain preamplifiers (HS-27, Neuralynx, Bozeman MT, USA). These signals were then digitized with 24-bit analog data acquisition cards with 30 dB onboard gain (PXI-4498, National Instruments, Austin, TX) and sampled at 32 kHz. Broadband signals (0.5 Hz – 16 kHz) were continuously recorded using custom-built LabVIEW software for the duration of the experiment. For tetrode array data, the spike detection and spike sorting methods have been described previously (Ecker et al. 2014; Tolias et al. 2007). For the Utah array, spikes were detected from individual electrodes following the same procedure. In this study, the term ‘multiunit’ refers to the set of all the spikes detected from a single tetrode or a single electrode (Utah array).

### Behavioral task

Visual stimuli were presented in a dark room using dedicated graphics workstations using Psychophysics Toolbox 3 (Brainard 1997; Kleiner 2007; Pelli 1997). For all experiments with monkeys A, CH and CL we presented stimuli on CRT monitors (model: Sgi C220 Flat Diamondtron; display size: 22 × 16° from a distance of 100 cm; resolution: 1600 × 1200 pixels; refresh rate: 100 Hz). For monkey L, we presented stimuli on an LCD monitor (Samsung – model S23A950D; refresh rate of 120 Hz; monitor resolution: 1920 × 1080 pixels, subtending visual angles of 29 × 16° from a viewing distance of 100 cm). We gamma-corrected the monitors to achieve a linear luminance response profile. The monitor background luminance was 6.1 cd/m^2^ (monkeys CL & A), 9.5 cd/m^2^ (monkey CH) or 0.04 cd/m^2^ (monkey L). The monkeys sat in a custom primate chair at 100 or 107 cm from the stimulus display monitor. Eye positions were continuously monitored online with scleral search coil for monkeys A, CH and CL and using a custom-built video tracker (frame rate: 250Hz) for monkey L. Eye position signals were also saved for offline analysis. Each trial (Fig. 4A) began with a brief sound that instructed the monkeys to start fixating at a red dot (0.12–0.14°) within a circular window of radius of 0.5–0.6° of visual angle. After the monkeys fixated for 300 ms, we presented different visual stimuli. The monkeys fixated for an additional 300 ms after the stimulus offset. For successfully completing the trials, the monkeys received juice or water reward. The next trial began after an inter-trial time of 1500 ms.

### Receptive field mapping

We mapped the spatiotemporal receptive fields using a white noise random dot stimulus. On a gray background, we presented black and white squares (0.11–0.14° side) on a rectangular grid covering the receptive field of all recorded neurons. The squares were presented one at a time for three video frames (25–30 ms) in a pseudorandom sequence for 1200–2000 ms. The sequence consisted of many iterations, in each of which every grid location was visited exactly once in a random order, thus balancing the number of times each location was visited over the course of the experiment. The monkeys performed 242 ± 56 trials (mean ± S.D.) in a session that lasted for around 20 min. Since primate V1 contains many complex cells and we were interested primarily in the location of the receptive fields, we performed reverse correlation ignoring the sign of the stimulus (i.e. both black and white were treated as positive). We assessed the quality of the receptive field estimation by the following heuristic method. We first averaged the receptive field maps obtained at lags ranging from 40 to 100 ms, resulting in a single spatial kernel for each multiunit. We fitted the spatial kernel with a two-dimensional Gaussian and computed the percentage of variance explained (across pixels) by the model. For all analyses in this study, we included multiunits for which the model explained more than 75% of the variance. From the model fitting, we also extracted receptive field centers and outlines. For illustration we outlined receptive fields by the elliptical contour at two standard deviations from the center.

### Speed manipulation experiment

Monkeys A, CH and CL were used in this experiment. Moving and flashed vertical bars of identical luminance and size (0.28 × 1.7°) were used as visual stimuli. The bar luminance was either 23 cd/m^2^ (monkeys A & CL) or 37 cd/m^2^ (monkey CH). We defined a stimulus presentation center for each monkey as the average of the receptive field centers (ARFC) of the neurons we recorded from; the mean eccentricity of this location was 1.5 ± 0.11° (azimuth: 0.87 ± 0.3° and elevation: 1.2 ± 0.3°; mean ± S.D.). In each stimulus period, only a flash or a moving bar was presented. We presented flashes for one video frame (10 ms). Since we recorded from many neurons simultaneously, to stimulate all the recorded neurons, we presented flashes at 5–7 locations around the ARFC (Fig. 4B). These locations were abutting each other without any overlap. The trajectory length of the moving bar was 4.6 or 5.4°. The midpoint of the moving bar’s trajectory was at the ARFC. The moving bar translated horizontally from left to right or from right to left at one of three speeds: 7, 14 or 28°/s (range: 6.9-7.4, 13.8-14.7 and 27.5-29.5 °/s respectively). All stimulus conditions were presented with equal probability. In each trial (Fig. 4A), we chose more than one stimulus condition randomly (two flashes and one moving stimulus for example) and presented them one after the other with an interstimulus period of 300 ms; this allowed us to use the monkeys’ fixating period efficiently and present multiple stimulus conditions within every trial. During the stimulus period of ≤ 1800 ms, we presented 4 ± 1 (mean ± S.D.) stimuli. In a session, we repeated each stimulus condition for 426 ± 216 (mean ± S.D.) times. The monkeys performed 1597 ± 718 (mean ± S.D.) trials per session. Each session lasted for 3 ± 1 (mean ± S.D.) hours.

### Luminance manipulation experiment

Monkey L was used in this experiment. The stimulus presentation followed the same overall design as the speed manipulation experiment (see above) with the following exceptions. The size of the bar was 0.15 × 1.8°. Moving and flashed bars with luminance values of 0.24, 0.82, 9.4, 48 cd/m^2^ were presented in each session. Flashes were presented at one of nine abutting locations with the ARFC at an eccentricity of 0.92 ± 0.07° (azimuth −0.46 ± 0° and elevation 0.79 ± 0.08°; mean ± S.D.). The trajectory length of the moving bar was 8.7°. The moving bar translated horizontally from left to right or from right to left at 18°/s. In the stimulus period of each trial, we presented 5 ± 1 (mean ± S.D.) stimuli. Each stimulus condition was repeated 120 ± 46 (mean ± S.D.) times. The monkey performed 1128 ± 432 (mean ± S.D.) trials per session with each session lasting 2 ± 1 (mean ± S.D.) hours. Note that to fit all luminance conditions within the recording duration, we did not test multiple speeds. Instead we chose a speed (18°/s) that was intermediate between speeds 7 and 28°/s that were used in the speed manipulation experiment. We also reduced the width of the bar to roughly half (0.15°) that of the bar used in the speed manipulation experiment (0.28°) so that when the bar moves, the footprints of the bars in the trajectory are contiguous without overlap or leaving a gap between adjacent instantaneous positions. The flash duration (8.3 ms) is also shorter than that used for speed manipulation (10ms) because we had to use an LCD monitor which had a higher refresh rate (120 Hz). We specifically chose an LCD monitor over the CRT monitor because to test very low luminance levels, we had to set the background luminance to lowest possible value; at that setting (but not at the background used in speed manipulation experiment), when the bar moved on the CRT monitor, it left behind a trail of phosphorescence that was obvious to a human observer. Such trailing luminance was not observed on the LCD monitor.

### Control experiment

Monkeys A and CL were used in this experiment. Stimuli were presented as outlined in the speed manipulation experiment. However, in addition to presenting flashed and moving bars separately as above, we also interleaved additional stimulus conditions where we presented the flash and moving bar together in two arrangements A1 and A2 (Fig.11A). In A1, we presented a flash inside the receptive fields and the moving bar below the flash but outside the receptive fields. To mimic the psychophysical experiment of the flash lag illusion, in arrangement A1, when the instantaneous position of the moving bar hit the azimuth of the ARFC, a flash was presented at one of 5–7 horizontal spatial offsets (0°, ±0.27°, ±0.55°, ±0.82°). We assigned a negative sign to the offsets if the flash appeared ahead of the moving bar along the motion direction and a positive sign if the flash appeared behind the moving bar. In arrangement A2, the vertical positions of the flash and moving bar in arrangement A1 were interchanged. The moving bar translated at a speed of 14°/s. The vertical center-to-center distance between the flash and the moving bar was 2.1°. With the bar height being 1.7°, the edge-to-edge gap between the flash and the moving bar was 0.4°. In each trial, we presented 3±1 (mean ± S.D.) stimulus conditions. Each stimulus condition was repeated 159±81 (mean ± S.D.) times. The monkeys completed 1930± 742 (mean ± S.D.) trials per session with each session lasting 3±1 (mean ± S.D.) hours.

### Electrophysiological dataset

For the entire study, we recorded neural data from a total of 1457 multiunits (monkey A: 288 CH: 191, CL: 306 and L: 672) over 62 sessions (A: 12, CH: 23, CL: 20 and L: 7) in an average period of six weeks from each monkey (A: 4, CH: 12, CL: 6 and L: 2). For the flash, relative to the pre-stimulus fixation period, majority (1038 (71%), A: 247, CH: 180, CL: 276 and L: 335) of the multiunits showed significantly enhanced responses measured over a window of 30–130 ms after the flash onset. A minority (44(3%), A: 2, CH: 11, CL: 20 and L: 11) of the multiunits showed flash-evoked suppression. For analyses, we included a subset of the multiunits (915 (63%), A: 237, CH: 166, CL: 256 and L: 256) that showed enhanced flash-evoked responses and passed the receptive-field-based selection criterion (955 (66%), A: 247, CH: 176, CL: 271 and L: 261, see *Receptive field mapping* section). After the above selections, one multiunit from monkey A was excluded from the analyses in Fig. 7 and Fig. 10 as its receptive field center was outside the flashed region. For the speed manipulation experiment, a total of 163 (A: 57, CH: 56 and CL: 50) single units were isolated out of which 44% (total: 71, A: 32, CH: 12 and CL: 27) met the selection criteria described above. For population decoding we chose all the single units from monkey CL since it had the most well-isolated units (median contamination measure (Tolias et al. 2007)(Interquartile range): CL: 0.039 (0.005, 0.086), CH: 0.076 (0.048, 0.117) and A: 0.092 (0.015, 0.142)).

### Response peak delay as neural representation delays for flash and moving stimuli

For the moving stimuli, assuming a receptive-field-based labeled-line code for position in V1, the latency of peak activity of a neuron closely approximates the representation delay. This is because, whenever there is a bar moving in the visual field, a population activity hill representing some moving bar position is simultaneously present in V1 (Fig. 2), except during the motion onset and offset. We assume that any subsequent visual area decoding moving bar position based on V1 activity would assign the instantaneous position of the bar center to the position encoded by the neurons whose activities maximally contribute to the peak of the hill. This would imply that the time at which a given neuron fires maximally is also the time at which the moving population hill activity is centered over this neuron’s topographic location in V1. Under this reasoning, the response peak latency would correspond to the latency of the V1 representation of the moving bar’s instantaneous position. For the flash, the situation is different because when a flash is presented in the visual field, a population activity hill starts to develop only after a delay. The hill then rises and falls over time without any change in the position of the peak of the hill. It is currently unknown at what point in time the activity hill fully represents the flash location. To be consistent with the method of latency computation of motion, we chose to compute peak response latency for flash as well.

### Estimation of flash response peak latency

For each flash condition, we first aligned the spike times of a given stimulus presentation to the flash onset time. We then computed mean firing rates across all stimulus presentations of a given condition after binning the spikes at half the monitor refresh period (4.2 or 5 ms). In each session, multiple flashes were presented, covering the receptive field of a given multiunit. We sought to find the mean firing rate response profile to a flash that was horizontally aligned with the center of the receptive field. However, there might not be any flash that was presented perfectly over the receptive field center since we did not optimize the flash locations for any particular neuron. In such cases, the mean firing rate profile that corresponds to a flash at the receptive field center was obtained by linearly interpolating the mean firing rate profiles of the flash locations left and right of the receptive field center. The mean firing rate response starting 150 ms before and ending 300 ms after the flash onset was then normalized (z-scored) to have zero mean and unit variance. After z-scoring, the responses of all multiunits under a given condition were averaged and smoothed using a Gaussian kernel with a standard deviation of 10 ms. Peak responses latencies were then computed from these averages. The responses of individual single and multiunits to flashed and moving bars were sometimes multimodal. Since we had a much larger multiunit dataset compared to single units, we chose to extract the latencies from responses averaged across multiunits. This procedure turned out to be more robust than extracting latency for each unit (for a description of how we estimated confidence intervals on the latencies, see section *Statistical Analysis* below).

### Estimation of motion response peak latency

For each motion condition, we aligned the spike times of a given presentation to the time at which the moving bar hit the center of the receptive field (i.e., the response time is set to zero when the moving bar’s instantaneous position matched the receptive field center). Since the moving bar occupied discrete positions along the trajectory that did not necessarily coincide with the receptive field center, we linearly interpolated the trajectory time points to obtain the time at which the trajectory crossed the receptive field center. We then computed mean firing rate across all presentations of a given condition after binning the spikes at half the monitor refresh period (4.2 or 5 ms). The mean firing rate response starting 150 ms before and ending 300 ms after the zero-time point was then normalized (z-scored) to have zero mean and unit variance. This normalized response was then averaged across multiunits. After this step, we followed the same procedure as for the flash responses described above and computed response peak latencies for each stimulus condition. The latencies were then averaged across the two motion directions.

### Latency estimation in control experiment

In the control experiment, we computed response peak latencies for flashes from arrangement A1 and for moving bars from arrangement A2 (see section *Control Experiment* above). To compute flash response latency for a given spatial offset, we first selected multiunits whose receptive field centers were within the spatial extent of the presented flash. Response peak latency was then extracted from this set of multiunits as described under the section *Estimation of flash response peak latency*. To compute the motion response latency for any spatial offset, we first selected multiunits whose receptive field centers were within the spatial extent of the moving bar when it hit the ARFC. Since the flashes were presented at different horizontal locations when the moving bar hit the ARFC, the same set of multiunits were used for extracting latencies under different spatial offsets. Motion response peak latencies were then computed as described under the section *Estimation of motion response peak latency*. Note that we chose to include a spatial offset for analysis only if there were more than ten multiunits for that condition. With this criterion, only the three spatial offsets around the ARFC qualified.

### Statistical analysis

All the statistical analyses on the neural data were done by bootstrapping (Efron and Tibshirani 1994). From the response (averaged over multiunits) peak latencies of flash and moving bar under various conditions, we computed the following test statistics: latency difference between flash and moving bar (Fig. 7B, Fig. 12-Fig. 13, F); slope of the trend in the latencies (Fig. 7B, Fig. 12-Fig. 13, F); latency differences and perceived spatial offset equivalents when changing speed (Fig. 7C & D, Fig. 10E & F, Fig. 12-Fig. 13, G & H) and luminance (Fig. 9C, F & G, Fig. 10H & I, Fig. 14G-I); and for the control experiments: latency differences across multiple spatial offsets (Fig.11B); latency differences for stimuli presented in isolation versus in combined condition (flash and moving bar presented together) (Fig.11C). To obtain significance levels and confidence intervals on these test statistics, we repeated 2000 times the entire procedure that generated a test statistic, each time with a different random set of multiunits obtained by resampling with replacement. Since the electrodes were implanted chronically, individual recordings from different days may not represent independent samples. To ensure that we use only independent samples for bootstrapping, we sampled electrode identities and included all units obtained from sampled electrodes. This procedure estimates the unit-to-unit variability without being confounded by dependent samples due to chronic recordings. From this bootstrap distribution, we computed the 95% percentile confidence intervals, which are reported as error bars. We defined the significance level (p-value) as *p* = 2 min(*q*, 1 − *q*), where *q* is the percentile of zero under the bootstrap distribution (this analysis assumes that the bootstrap distribution is an appropriate measure of the variability under the null hypothesis).

### Human psychophysics: task

Two human subjects (authors MS and SP) performed the standard flash lag psychophysical experiment as described previously (Subramaniyan et al. 2013). The subjects sat in a dark room with their heads stabilized by a chin-rest. After the subjects dark-adapted their eyes for five minutes, the stimulus presentation began. The subjects were simply instructed to stay fixated at the fixation spot during stimulus presentation; their eye movements were not tracked. In any given trial, we presented a flash below another bar that moved from left to right; the gap between the bottom edge of the moving bar and top edge of the flashed bar was 0.3° and both bars had identical luminance. We used seven different horizontal offsets between the flash and moving bar centers. The offset values ranged from around −6.3° to 2.4° in steps of around 1.5°. We used a constant flash location and created the spatial offsets by choosing the time of flash relative to the instantaneous position of the moving bar. To be comparable to the physiological experiments with monkey L, we made sure that at zero spatial offset, the average position of the flash and moving bar centers matched the ARFC used for monkey L. In each session, we randomly interleaved four bar luminance values. These luminance values, bar dimensions, monitor background luminance and speed of the moving bar were identical to those used in the luminance modulation experiment with monkey L, although here a longer motion trajectory of 18° was used. Using a keyboard, the subjects reported if the leading edge of the moving bar was on the right or left side of the flash at the moment the flash appeared. The subjects completed a total of seven sessions (MS: 5, SP: 2). In most sessions, we presented a total of 28 stimulus conditions (7 offsets × 4 luminance values × 1 motion direction × 1 speed). Each condition was repeated 20 times giving about 560 trials per session. Each session lasted for an average 23 min.

### Estimation of perceived spatial offset

To quantify the perceived spatial offset, we first converted the subjects’ responses into a probability of reporting that the moving bar was ahead of the flash. Then we fitted a logistic function to these probabilities as a function of spatial offsets, using psignifit3.0 toolbox (Frund et al. 2011; Wichmann and Hill 2001a; b). In the toolbox, we chose the constrained maximum likelihood method for parameter estimation and parametric bootstrapping for estimation of confidence intervals for parameters. We constrained the upper and lower asymptotes of the psychometric function to be equal with the prior distribution being a uniform distribution on the interval [0 0.1]. We defined the perceived spatial offset as the point of subjective equality, that is the veridical spatial offset at which subjects reported that the moving bar was ahead or behind the flashed bar with equal probability. To examine how the perceived spatial offset changed with luminance, we pooled the responses across sessions for each bar luminance before fitting the psychometric function. To perform statistical tests however, we fitted psychometric function for each session separately and computed perceived spatial offset.

### Statistical analysis of psychophysical data

For all statistical test on psychophysical data, linear mixed models were constructed in the statistical software PASW-18, with the following common settings: subjects were treated as random effects and perceived spatial offset as dependent variable. Specifically, the slope of the trend of the perceived spatial offset as a function of bar luminance (Fig. 9E) was tested for significance using the bar luminance as a covariate with the session start times set to indicate repeated measures. To test the effect of motion condition (foveopetal versus foveofugal) and speed on the perceived lag (Fig. 10B & D), speed was used as a covariate and motion condition as a factor, with the combination of motion condition and session start times set to indicate repeated measures.

### Probabilistic population decoding

The decoding method used here was chosen for its simplicity and its suitability for our experimental conditions abstracting away from neuronal implementation level details. Our goal was to decode the stimulus position presented to the animal from the single- or multiunit population activity based on the framework of probabilistic population coding (Dayan and Abbott 2005; Ma et al. 2006; Zhang et al. 1998). We took advantage of the fact that the motion stimulus we used was essentially a sequence of flashes. Hence, to decode the moving bar location, we first model the spatial encoding by measuring the population activity for the different flashed locations of the bar. Then, when the moving bar was presented, we decoded its instantaneous position by identifying the bar stimulus that was most likely, given the population activity at that instant. Note that in our experiments, only part of the motion trajectory overlaps with the space covered by the flashes. Since the spatial encoding is based on flash responses, we restricted the motion decoding to the region of the trajectory overlapping the flash locations. The decoding method is formalized as follows.

A flashed stimulus (Fig. 3A) evokes neural activity that extends over time outlasting the presence of the stimulus (~10ms) on the monitor (Fig. 3B). The post-stimulus period can be split into a sequence of contiguous time bins (of width Δt). We assume that, conditioned on the stimulus, the spiking responses are independent across both time and neurons. That is, activity (***R***) in any given time bin depends only on the stimulus location (S) and the elapsed time since stimulus onset (ε). Under this assumption, the neurons spike according to independent inhomogeneous Poisson distribution, with a time- and neuron-dependent mean spike count parameter *λ* (Fig. 3C). This produces the following probability distribution for neural activity (***R***) in a single time bin of width Δt:

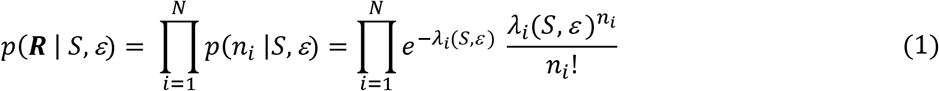

where

S - Stimulus (bar) at one of M possible locations (for example, see gray rectangles in Fig. 3A).
ε - Time elapsed since stimulus onset.
N – Number of neurons simultaneously recorded.
n_i_ - Spike count in a given time bin of width Δt for neuron *i*.
*λ_i_*(*S,ε*) – Mean spike count of neuron *i* in a time bin of width Δt after a delay of *ε* from stimulus (S) onset at one of M possible locations.
***R*** - (n_1_, n_2_,…,n_N_) - population activity (spike counts) in a single time bin.

**Fig. 3.**
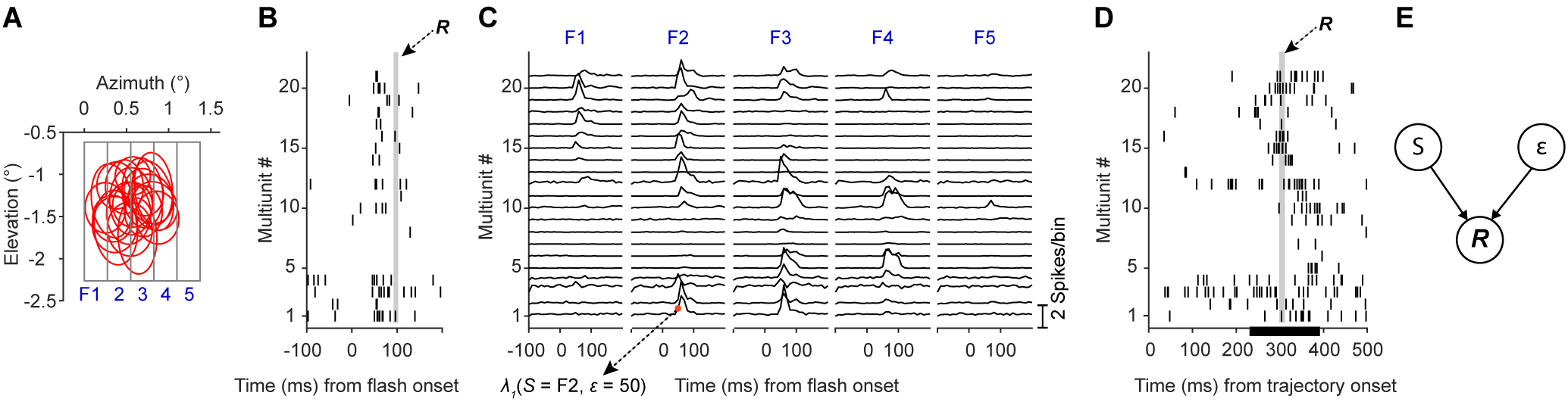
Probabilistic population decoding. ***A***, Outlines of receptive fields (red) of simultaneously recorded multiunits (n = 21) from a single representative session from monkey CL. The gray rectangles show the outlines of different flashes (labeled at F1, 2, 3, 4, 5) presented one at a time. ***B***, Single-trial raster plot of spiking responses (dark vertical bars) of all multiunits in ***A*** to a flash (F2). The spike counts within the thin gray vertical box (of width Δt =10ms) forms the activity vector (***R***) used in the decoding procedure. ***C***, Mean spike count across trials in 10ms consecutive time bins for all multiunits under each flash condition indicated on the top of each panel. An example value for *λ_i_*(*S,ε*) parameter (see main text) is indicated at the bottom left. The ordinate scale bar for the traces is shown on the bottom right corner. ***D***, Single-trial raster plot of spiking responses (dark vertical bars) of all multiunits in ***A*** to a bar moving from left to right at a speed of 7/s. Vertical thin gray rectangle as in ***B***. The black horizontal bar on the abscissa marks the time period the moving bar spent within the flashed region shown in ***A. E***, Graphical model of population activity. The population neural response at a given time bin (***R***) is governed by the stimulus (S) and the time elapsed (ε) since stimulus onset.

Note that, for the flash-evoked neural activity in any given time bin, the experimenter knows which flash stimulus caused the activity and how much time has elapsed since the stimulus onset (Fig. 3B). However, these two parameters are unknown from the brain’s perspective. In the case of the moving stimulus for which *a priori* we do not know the response latency, even the experimenter cannot know which stimulus location causes neural activity in a given time bin (Fig. 3D). This is due to the moving bar changing its location in every time bin leading to essentially multiple stimulus locations driving the neural activity in different time bins. As the experimenter cannot know which stimulus location caused the activity, he/she also cannot know how much time has elapsed since the onset of the stimulus (at a given location) driving the activity. For these reasons, in our decoding of flashed and moving stimuli, we treat the stimulus location and time elapsed since stimulus onset/arrival at a given location as random variables that follow a uniform distribution with flat priors. Note that the response at a single time bin for a moving stimulus is likely driven by *multiple* stimulus (bar) locations (spatiotemporal integration). However, to decode this activity, we are using an encoding model where population activity at each time arises from *single* stimulus (flash) locations. Hence in our decoding procedure, we are only approximating the spatiotemporal integration involved in generating population activity during motion. This leads to a graphical model (Fig. 3E) which, in combination with *Eq*.1, can be used to decode the stimulus position from the neural activity. Decoding this way in small time bins (~10ms) implies that a rate code is used by the brain for computing stimulus position. To compute the probability of a stimulus given the population activity in a single time bin, we first derive a joint distribution based on the model in Fig. 3E.

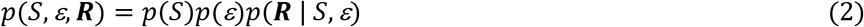

We assumed S and ε follow a uniform distribution (range of S: horizontal extent of flashed region, range of ε: 10 to ~175ms) and hence *p*(*S*) and *p*(*ε*) are constants (flat priors). We can then marginalize the above joint distribution over the elapsed time ε to compute the probability of a stimulus location given the population activity ***R*** in any arbitrary time bin:

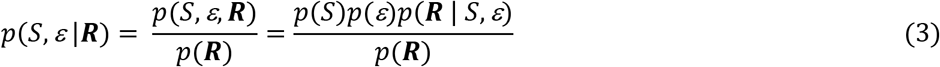

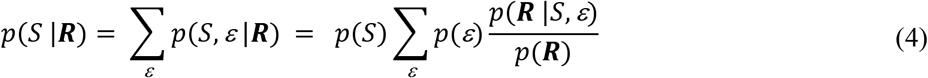

As *p*(*S*) and *p*(*ε*) are assumed to be constants for all values of *S* and *ε* respectively, they can be absorbed along with *p*(***R***) into the normalization constant *Z*(***R***) simplifying *Eq*.4 as:

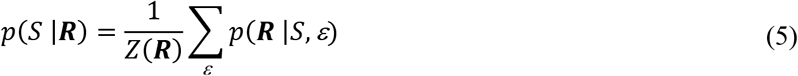

where *Z*(***R***) can be computed using the following normalization constraint:

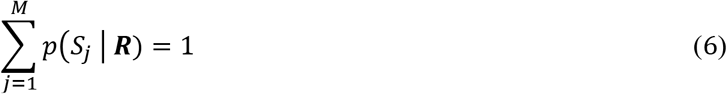

where the subscript *j* indexes the possible positions of the bar stimulus in visual space. Since the decoding was restricted to the space occupied by the flashes, the above constraint (*Eq*. 6) is justified as the decoded position should be within the flashed space. By using the above constraint, we avoided computing *p*(***R***) explicitly, as done by previous studies in similar decoding problems (Sanger 1996; Zhang et al. 1998). Note that for monkeys A and CH, although 7 flashes were presented in the task, we only included the central 5 flashes in the analysis as the flashes at the periphery did not have sufficient receptive field coverage.

Decoding was done trial by trial for each recording session using neurons recorded simultaneously. The same number of trials was used for all stimulus conditions within a session. In each trial, from stimulus onset, we stepped forward in small contiguous (non-overlapping) time bins (Δt = monitor refresh period, 8.3 ms for monkey L, 10 ms for others) and computed the posterior probability of each of the possible stimulus positions (M in total), given the population activity at that time bin. Hence, at every time instant, for a given test stimulus (flash or moving bar), we get an M-element vector of posterior probabilities that sum up to 1. For all stimulus conditions, the posterior probability was assigned to the end of the time bins. For example, the probability computed in the [0, 10) ms time bin was assigned to t = 10. This ensures that probability is causally related to the population activity. Also, note that for the speed of 7 °/s, only every fifth moving bar center matched flash centers (Fig. 5A). Hence when computing *λ*_i_(*S, ε*), we interpolated the mean firing rate from the flash centers to all positions (M in total) that the moving bar center occupied (white dots in panel D of Fig. 12-Fig. 14). For simplicity, the same interpolated *λ*_i_(*S, ε*) was used for all speeds. For the luminance modulation experiment, a similar interpolation procedure was done and the decoding of bar stimuli of a given luminance was based on encoding obtained from responses of flashes of matching luminance.

For the marginalization in *Eq*.4, we chose a time window that covers the flash-evoked responses of all recorded neurons for all monkeys. Based on visual inspection of the neural responses, this window was set to 10-175ms for monkey L (to allow for longer response latencies at low luminance conditions, see Fig. 8) and 10-150 ms for all other monkeys (see Fig. 6). The results and conclusion based on decoding are not sensitive to the exact values of the above time windows. For example, shortening the above windows to 20-130ms and 20-100ms respectively does not change the results and conclusions presented. However, including some time bins in which the population activity is at the baseline level minimizes the “edge effect” where the decoder, when decoding baseline-level activity, assigns a relatively higher probability to stimulus locations at the periphery (“edge”) of the flashed region (see the decoding in the 0-50ms window in Fig. 14B). This effect arises because the edge regions often have relatively poor receptive field coverage in our dataset (see first and last gray rectangle in Fig. 14A). When a bar stimulus is presented here, it evokes a population response which is similar to the baseline activity (see Fig. 3C, stimulus F5). Consider a decoder that does not include any baseline-level time bins in the marginalization time window in *Eq*. 5. When this decoder decodes baseline-level activity from any stimulus condition, it will assign a higher posterior probability to the edge regions (edge effect) as the bar stimuli at the interior locations are unlikely to evoke such poor baseline-level activity. Instead, including some baseline bins (e.g. bins with ε = 10 - 30ms) in the marginalization time window minimizes this effect. This is because, these bins contribute appreciably to the likelihood term inside the summation operation in *Eq*. 5. Hence, for a given *S*, the total likelihood summed over 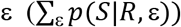 will be higher compared to when not including these bins. Moreover, as the baseline activity is similar for all bar locations (*S*), the large likelihood contribution will also be similar for all *S*. The result of such an overall increase in the total likelihood is that, after normalization in *Eq*. 5 (division by *Z*(***R***)), the posterior probabilities of the M locations become similar at times when there no stimulus evoked activity, thereby minimizing the edge effect.

### Cross-validation

The decoding was done on individual trials. Note that in the above model, we learnt the spatial encoding from the population response to flashes. Hence, when we decoded flash stimuli, to prevent overfitting, we kept aside a given trial for testing and used the remaining trials to train the model (i.e., compute the *λ*’s). This was then repeated for all available trials. For decoding the motion stimulus however, the separation into training and testing trials was unnecessary because the trials used for training (flash trials) were different from the trials in which testing was done (motion trials).

### Computing latencies from probabilistic decoding

The posterior probabilities of bar locations computed as described above were first averaged across trials and then across sessions for a given monkey. For estimating the decoding latency for flashes, we first averaged the probability values corresponding to positions within the horizontal spatial extent (see white horizontal bar in panel B top in Fig. 12-Fig. 14) of a flash stimulus. This was repeated for each flash condition and then the probabilities were averaged across the flashes (Fig. 12-Fig. 14, C) and smoothed with a Gaussian kernel with a standard deviation of ~5 ms. We then computed the latency of the peak of this averaged posterior probability as a measure of the latency of flash stimulus representation (Fig. 12-Fig. 14, F). For motion latency, first we aligned (centered) the probability vector computed at each time bin to the moving bar’s instantaneous position. We included time bins starting from the time the moving bar entered the flashed zone until about 120 ms or less after the bar exited the flashed region of space to account for latency of responses. The aligned vectors were then averaged (Fig. 12-Fig. 14 E) and smoothed with a Gaussian kernel with a standard deviation of 0.1°; the distance between the peak of the posterior probability and the origin, was taken as the spatial lag of the moving bar representation. The time delay (latency) corresponding to this spatial lag was then calculated by dividing the spatial lag by the speed of the moving bar. To obtain significance levels and confidence intervals on test statistics based on latencies computed as above, we repeated 2000 times the entire decoding procedure that generated a test statistic, each time with a different random set of single or multiunits within each session, by resampling with replacement.

## Results

We assessed if there are differences in the representation delays (latencies) of moving and flashed stimuli and whether this could account for the perceived spatial misalignment (offset) in the flash lag illusion. To this end, we recorded neural activity from V1 while the monkeys were shown either a flashed or a moving bar in a passive fixation task (Fig. 4A, see Methods). We performed two experiments: In the first, we varied the direction of motion and speed of the moving bar (7, 14 or 28 °/s), while keeping the moving and flash stimuli at a fixed luminance (23 or 36 cd/m^2^). In the second, we kept the speed constant and manipulated the luminance of the flash and the moving bar. In both cases, we measured the effect of the manipulation on the latency difference between the moving and the flashed bar and compared it to the psychophysical results from monkeys (Subramaniyan et al. 2013) and humans (Murakami 2001; Purushothaman et al. 1998; Subramaniyan et al. 2013; Whitney et al. 2000; Wojtach et al. 2008).

**Fig. 4.**
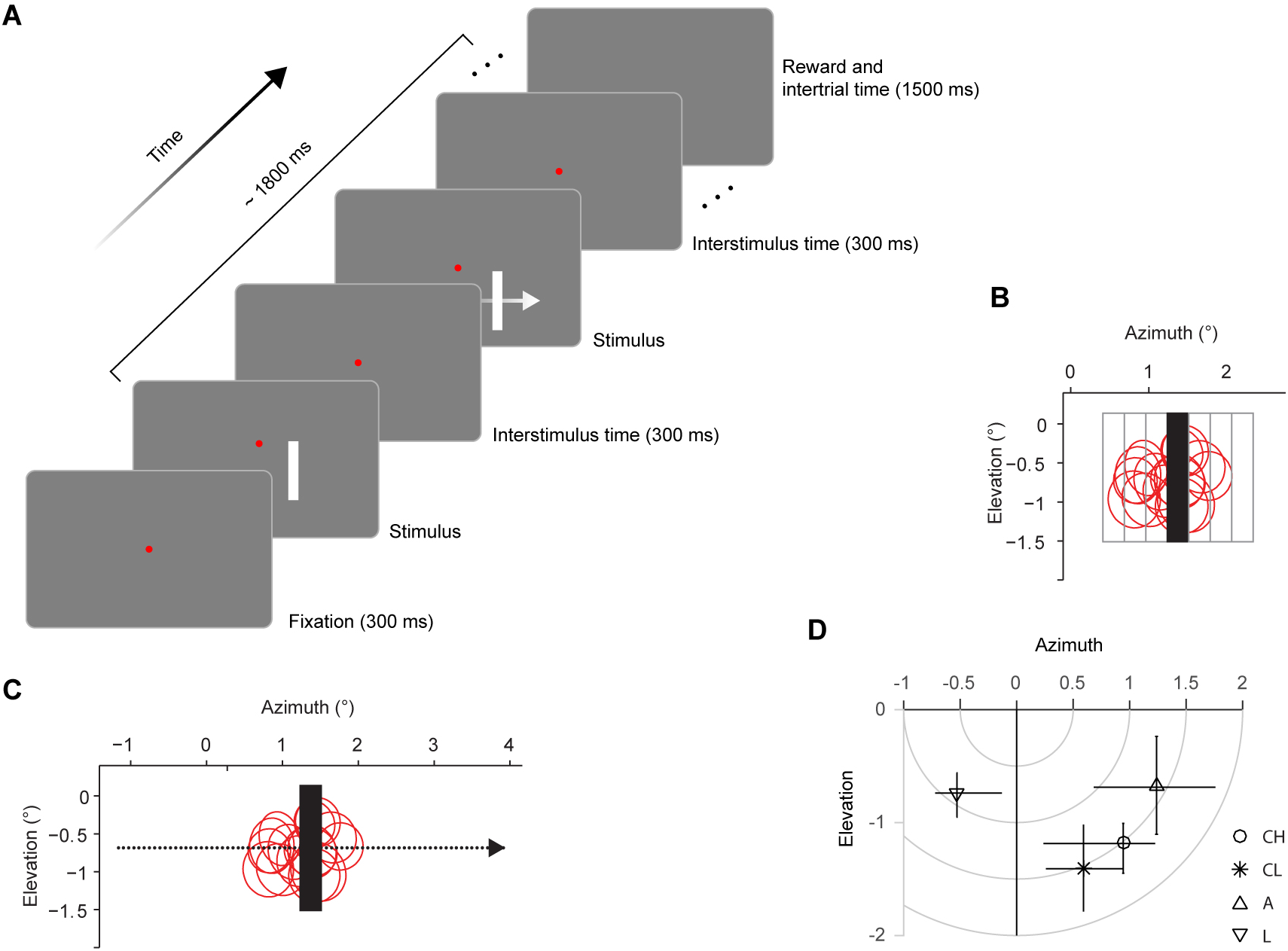
Fixation task and stimuli. ***A***, Monkeys fixated their gaze at a red circular dot at the center of the monitor within a fixation radius of 0.6°. After they maintained fixation for 300 ms, a single randomly chosen bright flash or moving stimulus was presented in a gray or dark background. The stimulus offset was followed by a 300 ms period in which no stimulus was presented except for the fixation spot. Then a randomly chosen flash or moving bar was presented again. With the monkeys maintaining fixation, this cycle continued until at most 1800 ms elapsed, after which they obtained a squirt of juice as reward. The next trial started after an inter-trial period of 1500 ms. ***B***, A flash (black bar) was presented at one of five to seven adjoining locations (gray rectangles) tiling the receptive fields (red circles) of all recorded neurons. ***C***, The moving bar (black bar) had the same size as the flash and moved from left to right or from right to left. The dots denote the positions of the bar center along the entire trajectory as the bar moved from left to right at a speed of 7 °/s. In ***B*** and ***C***, the coordinate (0°Azimuth, 0° Elevation) marks the center of fixation and the bars and receptive field outlines are drawn to scale. Note that the red circles show the outlines of only a subset of the recorded neurons. ***D***, Markers show median of receptive field centers of monkeys (CH, CL, A and L). The horizontal and vertical error bars indicate 95% percentile limits of azimuth and elevation respectively of receptive field centers. Isoeccentricity lines are shown in gray.

For experiment 1, we recorded from 523 multiunits in three animals using chronically implanted tetrode arrays. For experiment 2, we collected responses of 256 multiunits in one animal, using a 96-channel Utah array. After an initial receptive field mapping session, the main task began. We presented bright bars on a gray (experiment 1) or dark (experiment 2) background. In each trial either a flash or a moving bar was shown. Since we recorded from many neurons simultaneously, the flash locations were not optimized for any particular neuron. Instead, in each recording session, flashes were shown at five to seven fixed locations covering the receptive fields of all the recorded neurons (Fig. 4B). The moving bar swept across the receptive fields horizontally at a constant speed from left to right or from right to left with equal probability (Fig. 4C). For experiments 1 and 2, the receptive fields of units were in the right and left hemifield respectively (Fig. 4D). To test the predictions of different models of flash lag illusion, we estimated the stimulus representation delays of flashed and moving bars in V1 using two different approaches. The first method was based on the neuronal responses recorded on individual recording sites and the second one was based on decoding simultaneously recorded single- and multiunit population activity. Specifically, we tested the dependence of the latency difference between flashed and moving stimuli on speed, luminance and direction of motion.

### Dependence of latency difference on bar speed

We asked if the latency difference between the responses to flashed and moving bars depends on the speed of the moving bars. To this end, we recorded neural activity when a flash or a moving bar was presented and estimated response peak latencies using the receptive field (RF) center as a reference location (Fig. 5A). We then asked how long does the neuron take to reach its peak firing rate for a bar that is flashed at this location and for the same bar at the same location when it is part of a motion trajectory (Fig. 5A). For both stimuli, the time of response peak with respective to the time at which the bar appears (flash, Fig. 5B) or arrives (motion, Fig. 5C) at the same reference location, was taken as their respective representation delays (Fig. 5D). The assumptions behind using response peaks for computing representation delays are described in the Methods section. Note that when the speed increases, the time required for the bar to arrive at the reference location decreases (Fig. 5D). However, this difference in bar arrival times does not add to motion latencies as we measured latency after all stimuli arrive at a common reference location. The center of the receptive field is operationally defined as the region that elicits maximal response. Assuming an RF-based labeled-line code common for flashed and moving stimuli, if both stimuli are processed with the same delay, then, when either the moving or the flashed bar is at the RF center, they should both elicit their respective maximal response with the same delay. In other words, when the moving bar arrives at the RF center one would expect a peak response to occur with the same delay, because at that instant, the moving bar is indistinguishable from a flash. In contrast, we find that the response peak for all three moving stimulus conditions occurs earlier compared to that of flash (Fig. 5D). In addition, as the speed increases, the response peak latency also increases and approaches that of the flash. These observations suggest that a moving stimulus is processed differently from a flashed one and is represented earlier in time in a speed dependent manner compared to a flash in the same location.

**Fig. 5.**
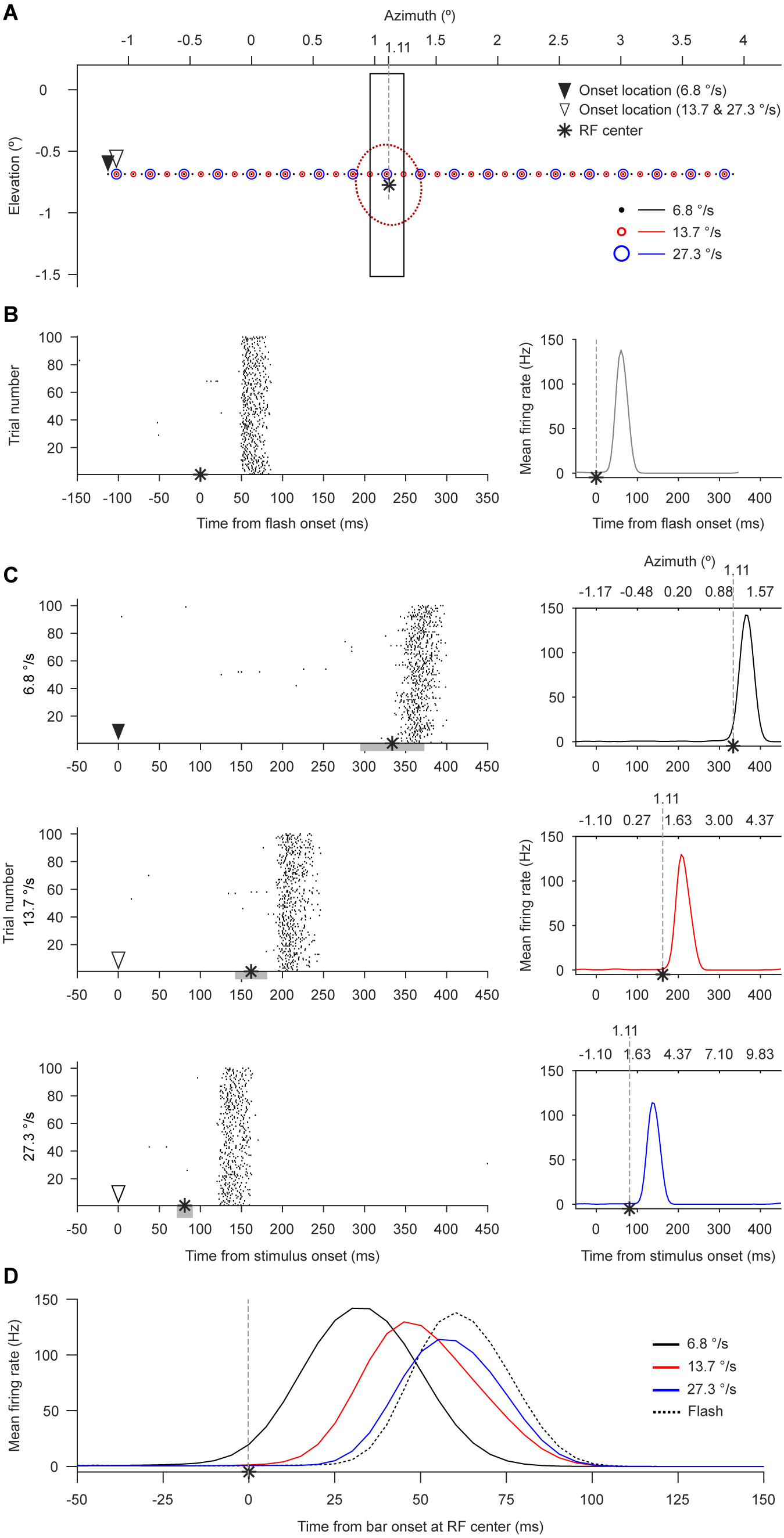
Single neuron responses to flash and moving bar and estimation of response peak latencies. ***A***, Illustration (drawn to scale) of the bar stimulus (rectangle) in visual space. The red dotted circle shows the 2-standard deviation outline of the neuron’s (from monkey A) receptive field (RF) with the RF center marked by the asterisk. In all panels of this figure, the vertical gray dashed line refers to the azimuth (1.11°) of RF center. In the flash condition, the bar is presented for one video frame as depicted. For moving conditions, the bar center occupies sequential positions marked by black dots (6.8 °/s), small red circles (13.7 °/s) or large blue circles (27.3 °/s); the bar shown is the instantaneous moving bar position that matches the flash. The triangles indicate the starting position of moving bar. ***B***, Left, raster plot showing neural responses to the flash shown in ***A***, aligned to stimulus onset time. Each dot denotes a spike and each row is a trial (only a subset of trials is shown). Right, mean firing rate plot for flash. ***C***, Left column, raster plots of responses for the bar moving from left to right at speeds indicated on the ordinate. Response times are aligned to motion trajectory onset time marked by the triangles. The gray horizontal bars mark the time needed to traverse the horizontal spatial extent of the receptive field outline shown in ***A***. Right column, mean firing rate plots corresponding to the respective raster plots shown on the left. In all subpanels of ***C***, the time at which the moving bar center crosses the receptive field center is marked by the asterisk. ***D***, Mean firing rate responses to all stimuli. Flash response is aligned to flash onset time. Moving bar responses are aligned to the time (asterisks in ***C***) at which the moving bar center crosses the receptive field center. The latency of response peaks for flash and moving bars is computed from this plot.

To estimate latency at the population level, we chose to first average the responses across the multiunits and then compute response peak latency from this average rather than vice versa. This was done because some multiunit responses had multiple response peaks, making it unclear as to which peak should be considered for latency estimation, and in experiment 2, the individual unit responses were too weak (Fig. 8) at the lowest luminance values to reliably find the response peak. Averaging the responses over the multiunits first, enabled us to robustly estimate latency and to apply a single procedure uniformly across all stimulus conditions.

Across our sample of multiunits from each monkey (Fig. 6), the peak response latencies for the motion condition at all three speeds were shorter compared to those for flashes (Fig. 7A & B; for each monkey, p < 0.0005, Bonferroni corrected, bootstrap test; see Methods). As the speed increased, the latency of the motion response approached that of the flash (Fig. 7B). Therefore, the latency difference between flash and motion decreased as the speed increased (Fig. 7C; p < 0.0005, bootstrap test) but remained greater than zero (p < 0.0005, Bonferroni corrected; bootstrap test).

**Fig. 6.**
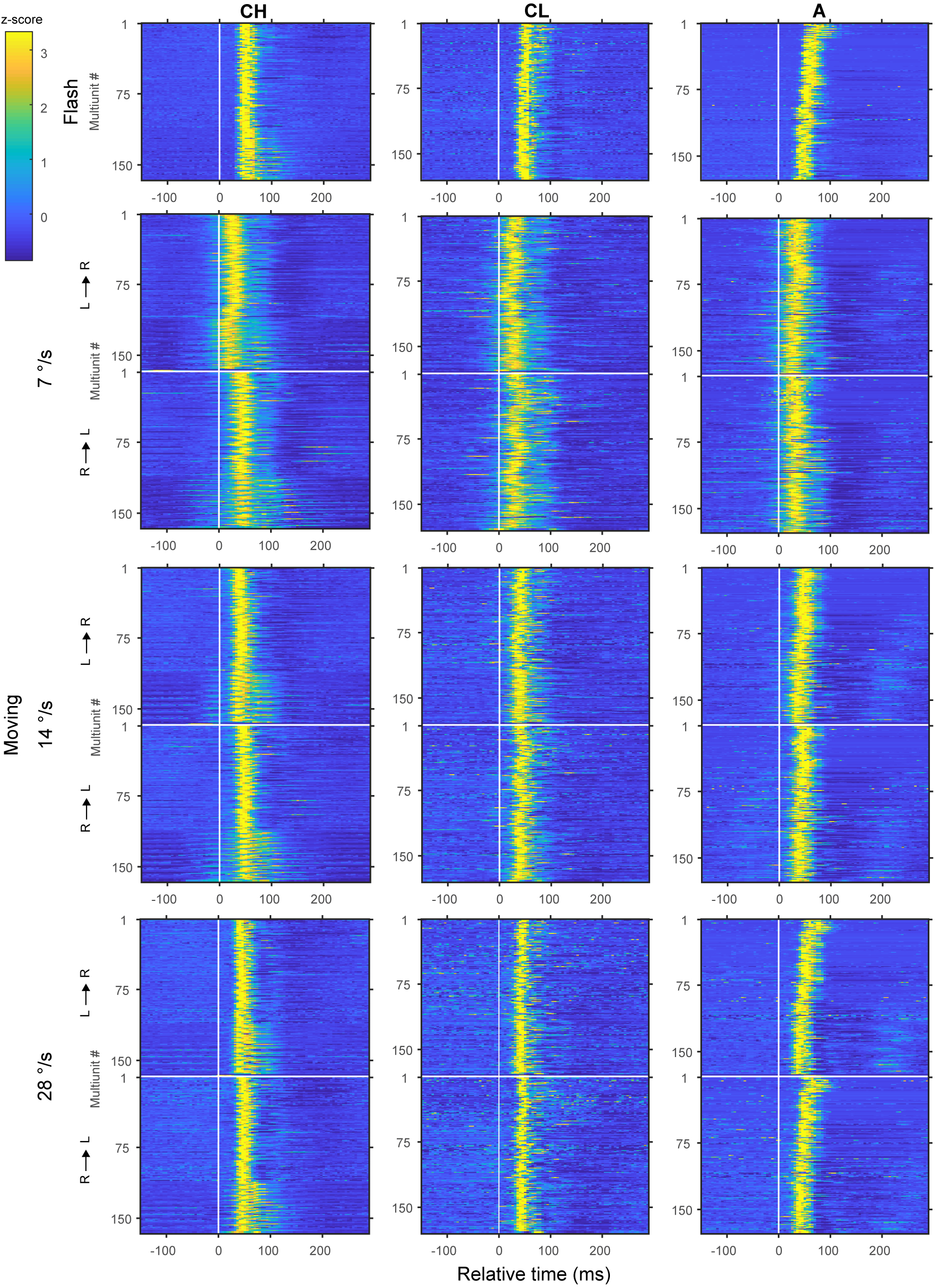
Trial-averaged responses of multiunits in the speed manipulation experiment in monkeys (CH, CL and A). Columns of panels represent monkey subjects indicated on the top. Rows of panels represent stimulus conditions indicated on the left. In each image, each row, ordered in ascending order of recording day, represents response of a multiunit. The vertical white line marks the time the stimulus hits the receptive field center. The horizontal white line separates the two motion direction conditions: L→R, motion from left to right; R→L, motion from right to left. Color range is clipped at 95^th^ percentile of responses grouped from all conditions.

**Fig. 7.**
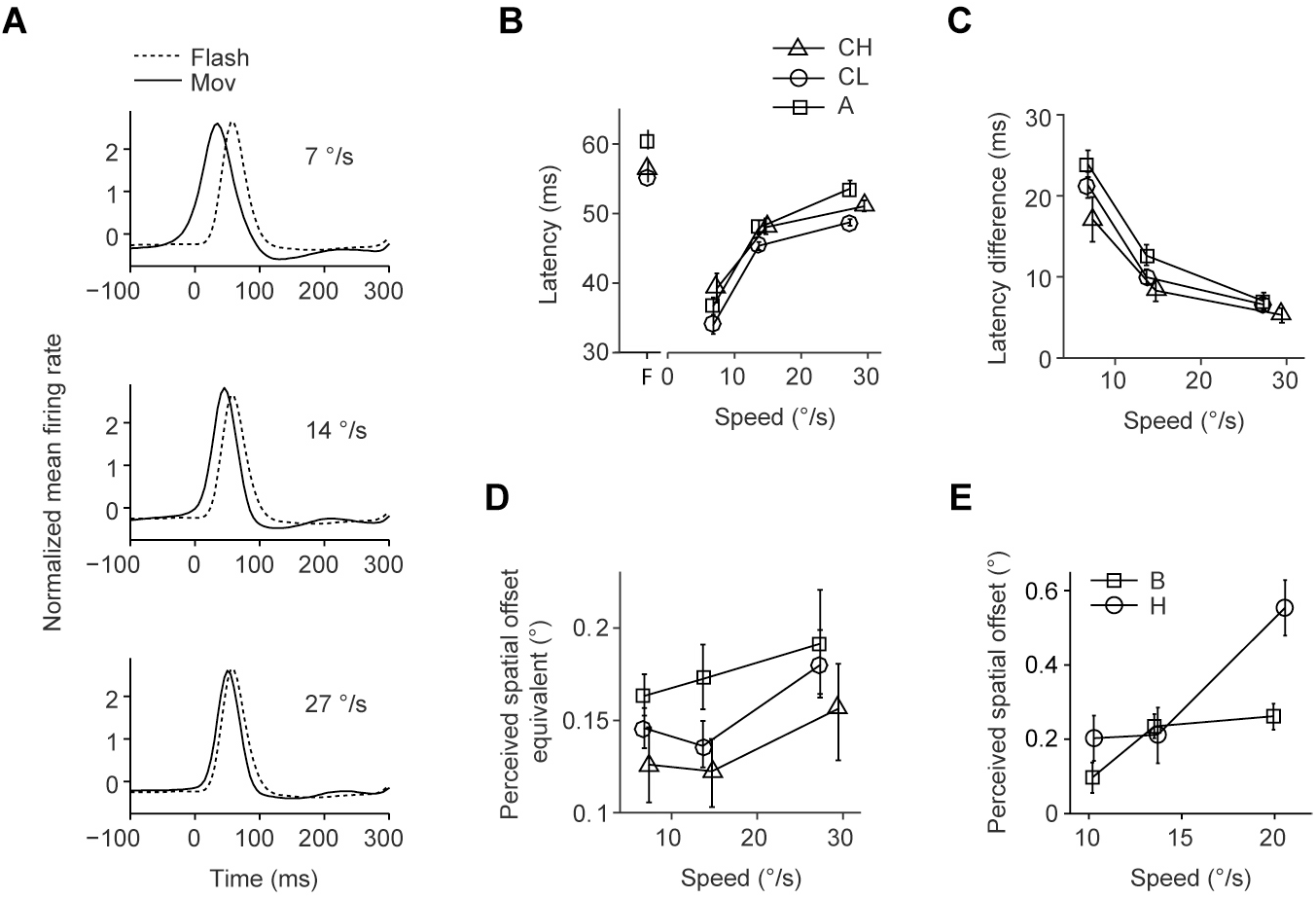
Population response and its correlation to flash lag psychophysics. ***A***, Normalized multiunit responses to flash (dotted trace) and motion (solid trace), averaged over all multiunits from monkey A (n = 177). The speed of motion is indicated on the top right corner of each panel. ***B***, Mean response peak latency for flash and motion plotted as a function of speed, for the three monkeys (n = 177, 166 and 180 for A, CH and CL respectively). The latencies for flash are plotted at the abscissa location marked by ‘F’. Error bars: 95% bootstrap percentile-based plug-in estimate of confidence intervals (note that most CIs are smaller than the markers). ***C***, Mean latency difference (flash latency minus motion latency) as a function of speed. Markers, sample size and error bars are as in ***B. D***, Speed dependence of perceived spatial offset equivalent computed from latency differences shown in ***C***. Markers and error bars are as in ***B. E***, Speed dependence of perceived spatial offset measured from two separate monkeys (B and H), re-plotted here from Subramaniyan et al. (2013).

This effect is consistent with the speed dependence of the magnitude of the perceived spatial offset observed in the psychophysical data collected in macaques (Subramaniyan et al. 2013). In our electrophysiological experiments, we manipulated the speed and measured the representation latencies of flash and moving bar rather than the perceived spatial offset which cannot be computed directly from the neural responses since it is a subjectively perceived quantity. In the psychophysical test, the subjects report the relative spatial offset between the flash and moving bar rather than how far the moving bar lags behind its own veridical location. Hence in computing the neural equivalent of perceived spatial offset, the latency of moving bar alone cannot be used - it is the difference (*L_d_*) in the representation delays of flash (*L_f_*) and moving bar (*L_m_*) that is needed. The neural equivalent of perceived spatial offset (*X*) was then computed by multiplying the speed (*v*) by the latency difference, i.e.,

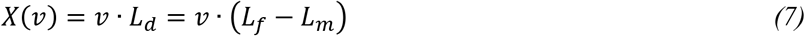

Although the latency difference decreased with speed, the perceived spatial offset equivalent increased with speed (Fig. 7D; p < 0.0005, bootstrap test). This counterintuitive effect can be explained by noting that the latency difference is not a constant but varies with speed (Fig. 7B). Hence,

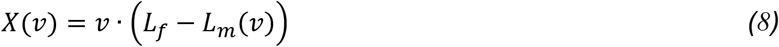

Differentiating both sides with respect to speed,

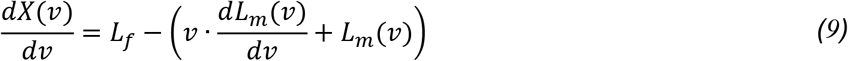

From *Eq*. 9, for the perceived spatial offset to increase with speed, i. e, for 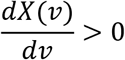,

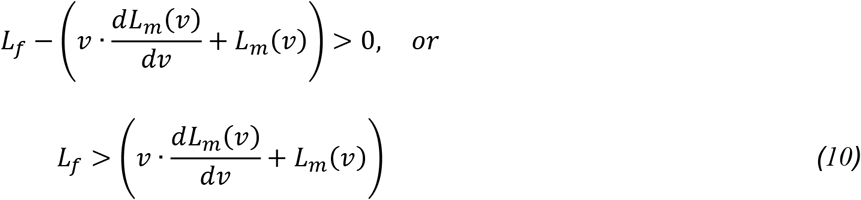

Hence, as long as *Eq.10* is satisfied, the perceived spatial offset will increase with speed even if motion latency increases 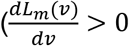, our data) or decreases 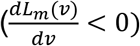 with speed.

The simplest case arises when motion latency does not change with speed 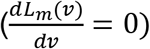, that is,

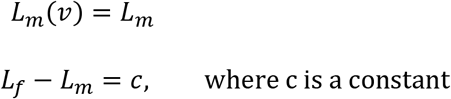

Therefore,

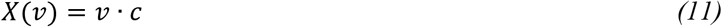

Hence *Eq.11* shows that perceived spatial offset linearly increases with speed as long as *L_f_* > *L_m_*. This assumption of constant motion latency is commonly made in psychophysical literature of flash lag illusion. Consistent with this assumption, some psychophysical studies have shown that the perceived spatial offset increases with speed (see Discussion). However, as indicated by *Eq.10*, this assumption is not necessary for explaining the increase in perceived spatial offset with speed. Our results demonstrate that despite an increase in motion latency with speed (Fig. 7B), the neural equivalent of perceived spatial offset increases with speed (Fig. 7D).

The increase of the perceived spatial offset equivalent with speed is consistent with our psychophysical results (Subramaniyan et al. 2013) from two other monkeys of the same species (Fig. 7E) and with human psychophysical studies (see Discussion). Together, these results show that in primary visual cortex, irrespective of the speed, the moving bar latency is not fully compensated (zero latency) as would be predicted by the spatial extrapolation model and that the latency of flash and moving bar are not equal as would be predicted by the motion-biasing model. On the other hand, our results are consistent with the differential latency model.

### Dependence of latency difference on bar luminance

The latency difference between moving and flashed bars may also depend on bar luminance. To test this, in the second experiment, we fixed the speed of the moving bar at 18 °/s and presented flashes and moving bars with luminance values of 0.2, 0.8, 9 and 48 cd/m^2^ (Fig. 8). We found that the motion response occurred earlier in time relative to the flash response (Fig. 9A). For all luminance values tested, the motion response peak latency was lower than that of the flash (Fig. 9B, p < 0.0005, Bonferroni corrected, bootstrap test). For both the flash and moving bar, the peak response latencies decreased as the luminance increased, although they decreased differently (Fig. 9B). Accordingly, the latency difference decreased as the luminance increased (Fig. 9C, p < 0.0005, bootstrap test).

**Fig. 8.**
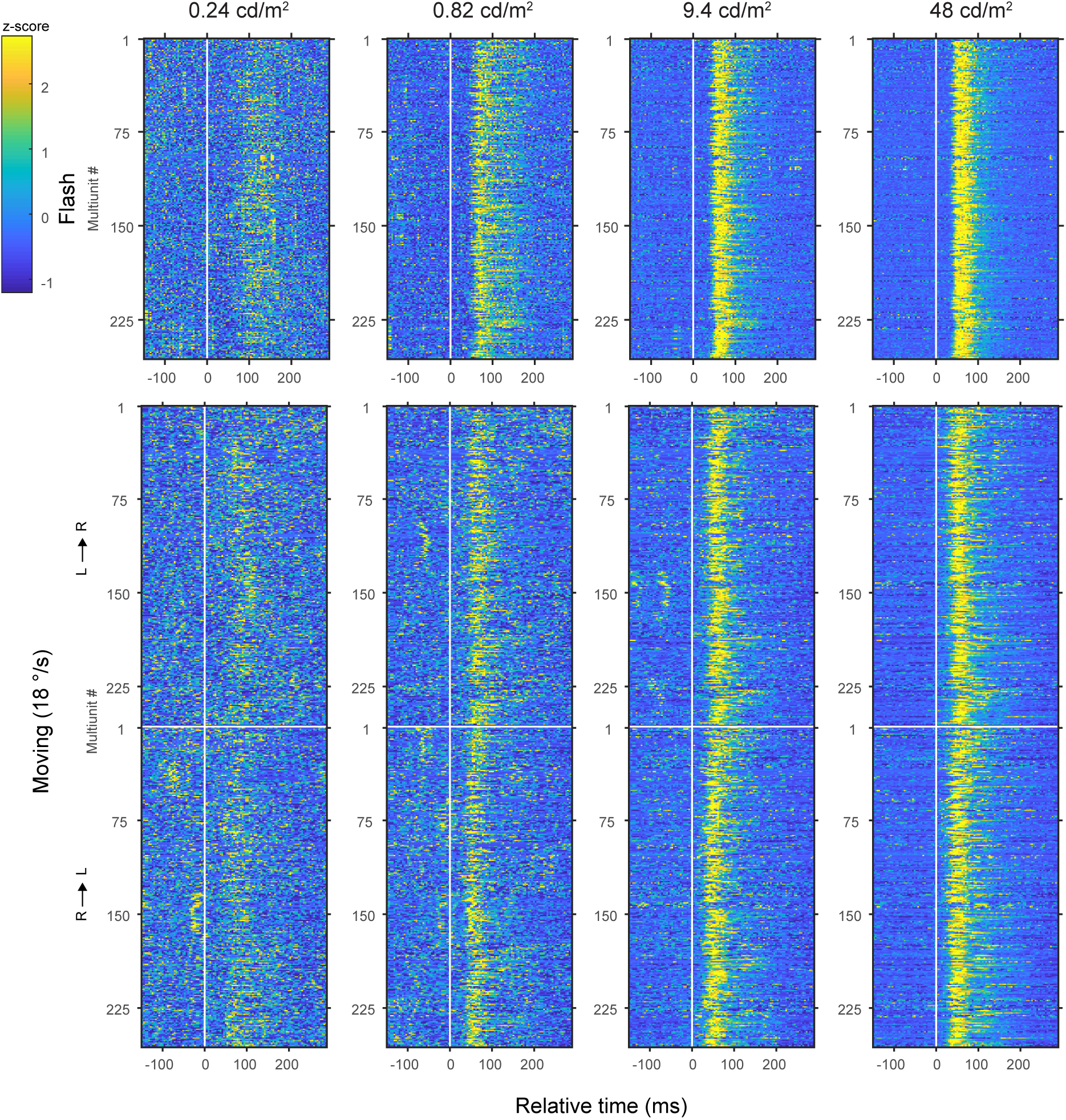
Trial-averaged responses of multiunits in the luminance manipulation experiment in monkey L. Columns of panels represent stimulus luminance indicated on the top. Rows of panels represent other stimulus conditions indicated on the left. In each image, each row, ordered in ascending order of recording day, represents response of a multiunit. The vertical white line marks the time the stimulus hits the receptive field center. The horizontal white line separates the two motion direction conditions: L→R, motion from left to right; R→L, motion from right to left. Color range is clipped at 95^th^ percentile of responses grouped from all conditions.

**Fig. 9.**
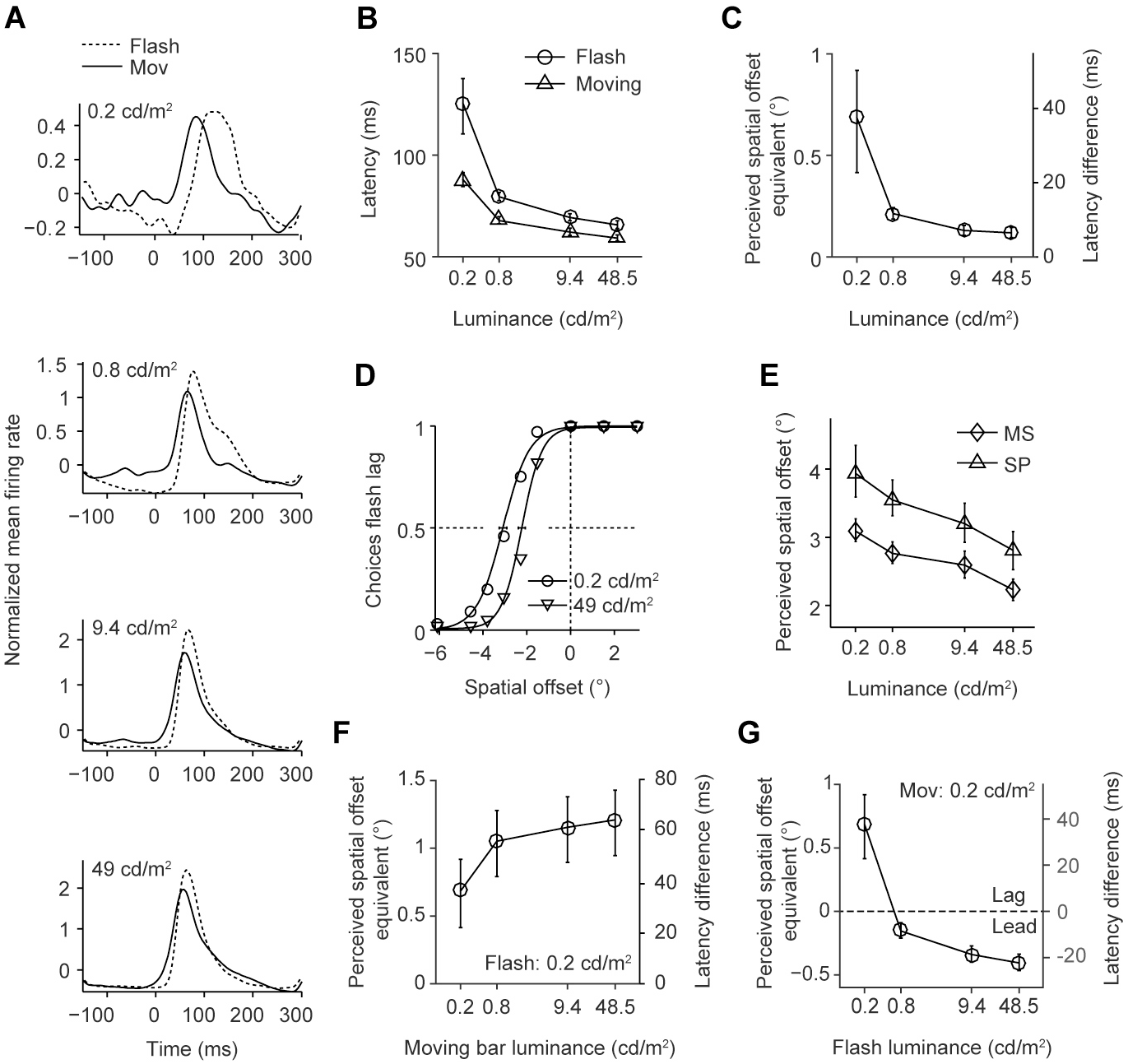
Luminance dependence of population response to flash and motion and its correlation to flash lag psychophysics. ***A***, Normalized firing rate responses to flash (dotted trace) and moving bar (solid trace) averaged across multiunits (n = 256) from all sessions (n = 7) from monkey L. In each subpanel the flash and moving bar had the same luminance (indicated on the top left corner). ***B***, Response peak latencies as a function of luminance, for flash and motion obtained from the data shown in ***A***. Error bars: 95% bootstrap percentile-based plug-in estimate of confidence intervals. ***C***, Luminance dependence of latency difference (flash latency minus motion latency, right vertical axis) computed from data shown in ***B***. Left vertical axis shows perceived spatial offset equivalent computed by multiplying the latency difference by speed (18 °/s). Error bars as in ***B. D***, Psychometric functions from human subject MS (for each data point, n = 100 trials, pooled from 5 sessions; for subject SP, n = 40 trials (2 sessions) per data point). The probability of the subject reporting that the flash is spatially lagging the moving bar is plotted against the veridical spatial offsets between the flash and moving bar at two luminance values (indicated at the bottom right corner). Error bars as in ***B. E***, Luminance dependence of perceived spatial offsets for human subjects (MS and SP). The perceived spatial offsets were computed from the psychometric functions using the method of compensation. Error bars as in ***B. F*** and ***G***, Latency difference and perceived spatial offset equivalent as a function of moving bar luminance (***F***) for a constant flash luminance (0.2 cd/m^2^) or as a function of flash luminance (***G***) for a constant moving bar luminance (0.2 cd/m^2^). The dotted line in ***G*** separates the luminance conditions that gave rise to perceived spatial offset equivalent corresponding to psychophysically measured flash-lag (‘Lag’) and flash-lead (‘Lead’) conditions. Error bars as in ***B***.

To compare physiological and psychophysical data, we again converted the latency differences into perceived spatial offset equivalent by multiplying the latency differences with speed (*Eq.8*). The perceived spatial offset equivalent decreased with luminance (Fig. 9C, p < 0.0005, bootstrap test). Although we currently do not have psychophysical data on the luminance dependence of the flash-lag effect in monkeys, we have previously shown that monkeys perceive the illusion similar to humans (Subramaniyan et al. 2013). We therefore measured perceived spatial offsets from two human subjects using the same luminance and stimulus parameters used for the monkey physiology. Indeed, the perceived spatial offset decreased with luminance in both observers (Fig. 9D and Fig. 9E; F (1, 24) = 14.6; p = 0.001; linear mixed model), in good agreement with the physiological results.

In the above analysis, we computed latency difference data between flash and moving bar with identical luminance and showed that they correlate well with human psychophysical data. Given that we presented each luminance condition in isolation, it is possible to compute the latency difference between a flash and a moving bar having different luminance values. In human psychophysics, when the flash luminance is fixed at a very low detectability level, the perceived spatial offset increases with the moving bar luminance (Öğmen et al. 2004; Purushothaman et al. 1998). To see if this is also evident in our neural data, we used the latency of the flash condition with the lowest luminance to compute latency difference at all moving bar luminance conditions. Interestingly, qualitatively similar to the human psychophysical results, we found that the perceived spatial offset equivalent increased (p < 0.0005, bootstrap test) with the moving bar luminance (Fig. 9F). An even more interesting psychophysical result is obtained when the moving bar luminance is fixed at a very low detectability level and the flash luminance in varied. For a sufficiently high flash luminance, the flash-lag effect is reversed where humans perceive the flash to be in front of the moving bar (flash-lead effect) (Öğmen et al. 2004; Purushothaman et al. 1998). We again found a qualitatively similar result in our neural data (Fig. 9G) where the perceived spatial offset equivalent decreased (p < 0.0005, bootstrap test) changing from being positive (flash-lag) to negative (flash-lead) as the flash luminance level was increased, correlating well with the human psychophysical results.

### Dependence of latency difference on motion direction

In addition to speed and luminance, the direction of motion has also been shown to affect the perceived spatial offset. Humans report a larger spatial offset for motion towards fovea (foveopetal, Fig. 10A) than motion away from fovea (foveofugal) (Kanai et al. 2004; Mateeff et al. 1991; Shi and Nijhawan 2008). We reproduced this finding in our stimulus paradigm where humans reported a higher spatial offset for foveopetal motion direction in a speed dependent manner (Fig. 10B, significant speed effect: F (1, 93.2) = 14.8, p < 0.001; nonsignificant motion condition effect: F (1, 75.8) = 2.56, p = 0.11; significant speed x motion condition interaction: F (1, 79.2) = 10.4, p < 0.01). Surprisingly, in the monkeys this motion effect was reversed under the same stimulus conditions (Fig. 10C & D, significant main effects and interaction: speed: F (1, 64) = 27.3, p < 0.001; motion condition: F (1, 67.6) = 12, p = 0.001; speed x motion condition: F (1, 64.8) = 6.6, p = 0.013). Correlating with this, the neural response latencies were lower (Fig. 10E) and the perceived spatial offset equivalent were higher (Fig. 10F & G), for the foveofugal condition in two of the three monkeys (latency and perceived spatial offset equivalent: p < 0.0005 for CH and CL and p > 0.05 for A; Bonferroni corrected for multiple speeds, bootstrap test). Note that in the neural data from all three monkeys (CH, CL & A), the receptive fields were in the right hemifield. Consequently, foveopetal condition is inseparable from motion from right to left visual hemifield and the neural effect we observed may reflect the later rather than the former condition. However, this is less likely for the following reason. In monkey L where we varied stimulus luminance, the receptive fields were in the hemifield opposite to that of the above other three monkeys (Fig. 4D). This led to the foveopetal condition being coupled with motion from left to right hemifield. Despite this, we observed the same effect found in the other data set (CH, CL and A), i.e., the latencies were lower (Fig. 10H) the perceived spatial offset equivalents were higher (Fig. 10I, p < 0.0005, Bonferroni corrected, bootstrap test) for foveofugal condition under all luminance values tested, suggesting that in monkeys, motion away from fovea produces a larger flash lag effect. The internal consistency between psychophysical and neural data within the monkey species strongly suggests that latency difference can explain a species-specific aspect of the flash lag illusion.

**Fig. 10.**
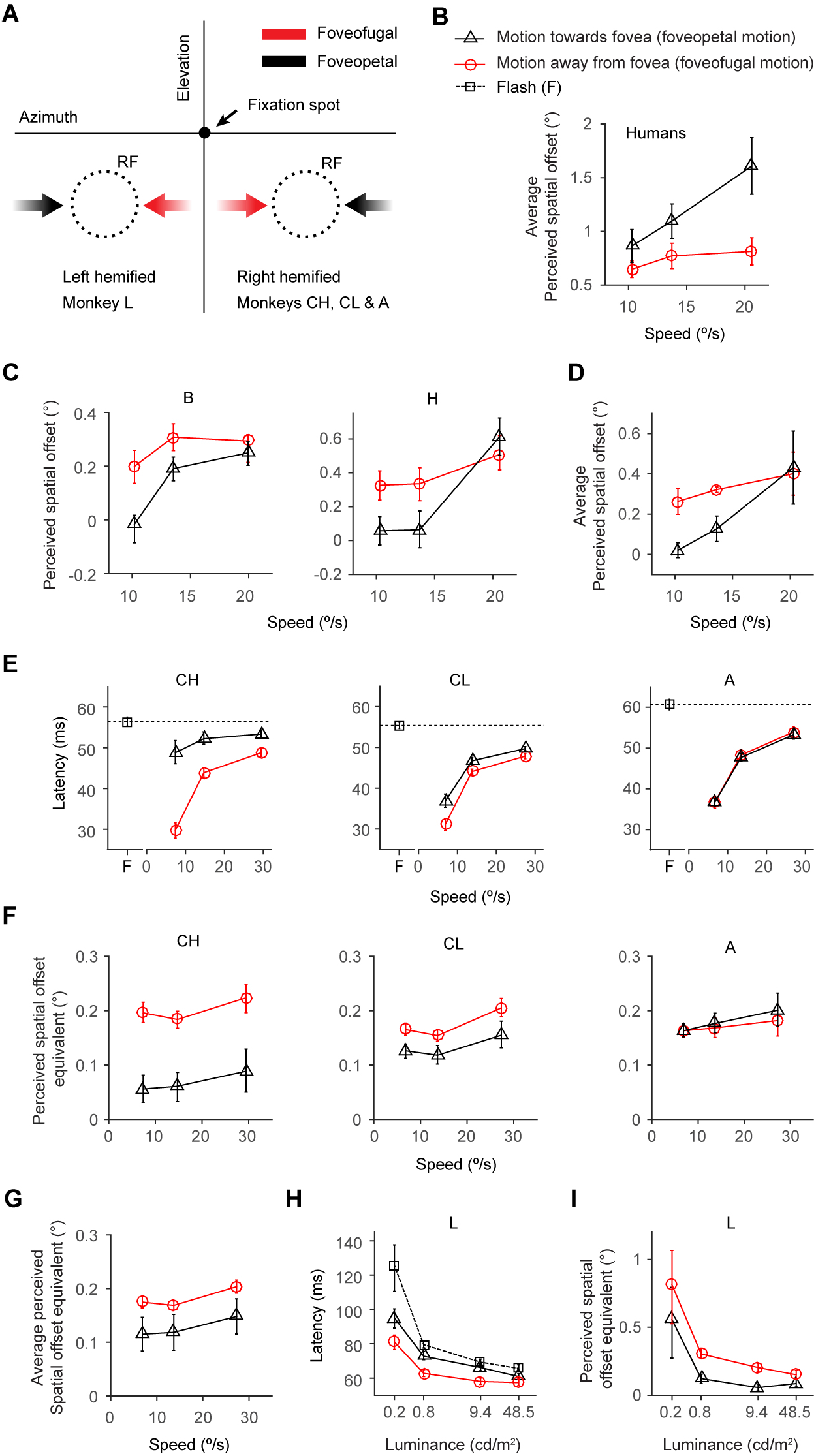
Effect of motion direction on perceived spatial offset and its neural equivalent. Illustration of motion directions (***A***) defined for a given neuron as *foveopetal* if the trajectory hits the receptive field (RF, dotted circles, hypothetical) before crossing the vertical meridian and *foveofugal* if the trajectory crosses the vertical meridian before hitting the receptive field. For psychophysics, the same convention applies with the subjects making relative position judgement of bar stimuli at the RF locations. Speed and moving direction dependence of average perceived spatial offsets in humans (***B***, n = 8) and monkeys (***D***, n = 2; individual monkey (B and H) data in ***C***) computed from data presented in (Subramaniyan et al. 2013). Speed and moving direction dependence of multiunit response peak latencies (***E***) and the perceived spatial offset equivalent (***F***) for the individual monkeys (n = 177 (A), 166 (CH) and 180 (CL)) and its average (***G***). Luminance and motion direction dependence of multiunit response peak latencies (***H***) and the perceived spatial offset equivalent (***I***) in monkey L (n = 256).

### Simultaneous presentation of flashed and moving stimuli

In summary, our physiological data from speed and luminance manipulation are in good agreement with psychophysical results and the predictions of the differential latency model of the flash lag effect. One potential caveat is that in our physiology experiments we presented the flashes and moving bars in isolation. However, to generate the flash lag illusion, the flashed and the moving bar are presented simultaneously with perfect alignment. It is thus conceivable that if we had presented the flash and the moving bar together, the results might have been different. To rule out this possibility, we conducted a control experiment in which we presented the flash and moving bar together at different spatial offsets, including a zero-offset condition where the flashed and the moving bar were in alignment. This allowed us to determine whether there is a change in latency as a function of spatial offset for simultaneously displayed stimuli.

We presented the flashes and moving bars simultaneously (‘combined’ condition) in two different arrangements. In the first, we presented flashes at the receptive fields and the moving bar (speed: 14 °/s) outside the receptive fields (Fig.11A, left panel) and vice versa in the second (Fig.11A, right panel), at 5-7 different spatial offsets in a gray background. For analysis, we chose the central three offset conditions that had sufficient number of multiunits (see Methods). We then computed the flash response peak latencies from the first arrangement and the motion response peak latencies from the second. The latency difference was not significantly different among the three spatial offsets (p > 0.76, bootstrap test). In the same recording sessions, we also presented flashes and moving bars in isolation inside the receptive fields. To test whether in the combined condition, a second stimulus affected response latencies, we pooled the latency difference data across monkeys and spatial offsets in the combined condition and compared it to those obtained where stimuli were presented in isolation (‘single’ condition; ‘s’ in Fig.11B & C). We found no significant difference between the combined and the single condition (p > 0.99, bootstrap test). These results suggest that in awake fixating macaques, the latencies of the flash or moving bar representation in V1 are not influenced by the presence of a second bar stimulus outside the classical receptive field.

**Fig. 11.**
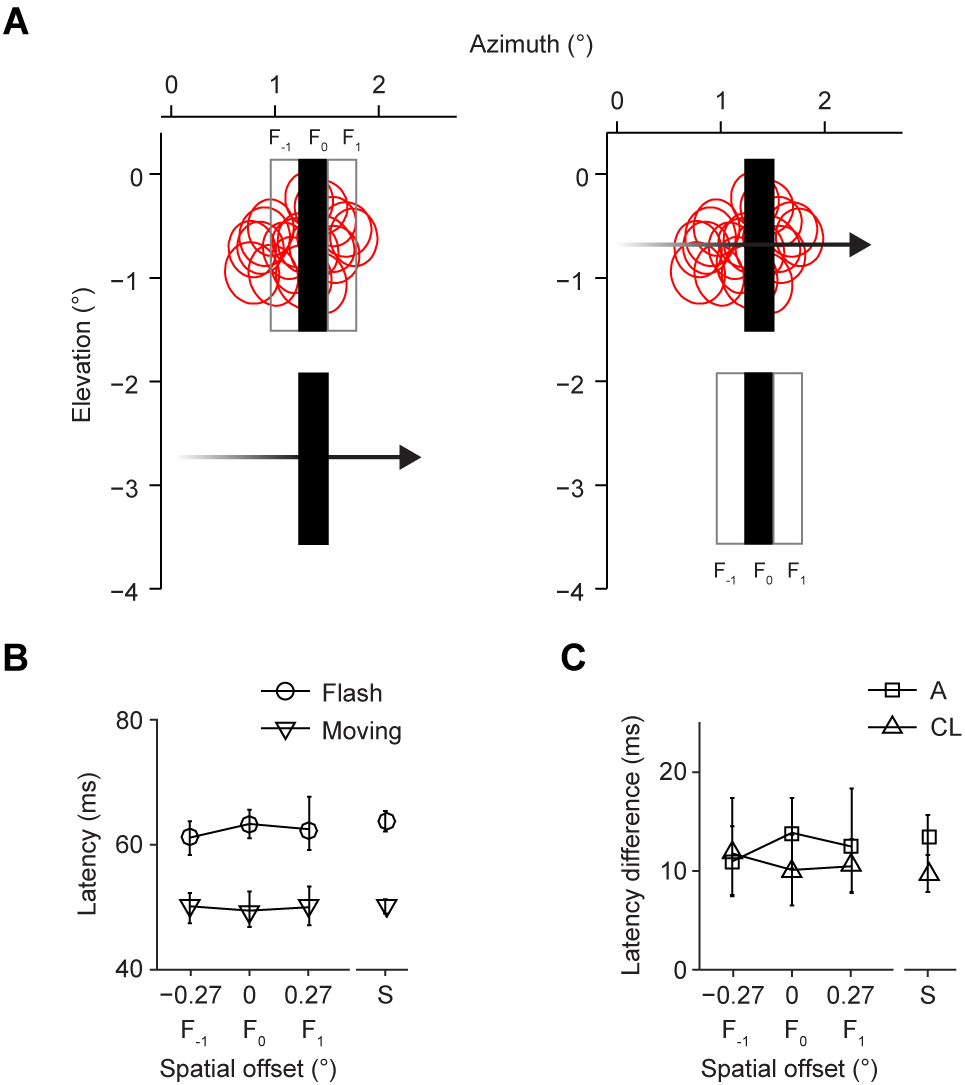
Control experiment. ***A***, Stimulus configurations. The red circles show the outline of receptive fields of a subset of the multiunits used in the analysis. Left panel: The filled rectangles show an example stimulus configuration with zero spatial offset. The two outlined rectangles show the other locations where we presented the flashes. Right panel: Same as the left panel except that the flash is now presented outside the receptive fields. Letter labels (F_−1_, F_0_, F_1_) identify flash locations whose relative horizontal offsets from moving bar form the abscissa in ***B*** & ***C. B***, Response peak latencies of flash and moving bar conditions from monkey CL (n = 25 ±7 multiunits per condition; mean ±1 S.D; for monkey A, n = 14 ±3 per condition), plotted as a function of the horizontal spatial separation between the flash and the instantaneous position of the moving bar (spatial offset). Data points from either flash or motion condition presented without an accompanying stimulus, are plotted at the abscissa location marked by ‘S’ (n = 76 multiunits; for monkey A, n = 59). Error bars: 95% bootstrap percentile-based plug-in estimate of confidence intervals. ***C***, Latency difference between flash and moving bar conditions from monkey A and CL, plotted as a function of spatial offset. Error bars as in ***B***.

### Population decoding of flashed and moving bars

The conclusions reached so far were based on latencies estimated by aligning individual neuronal responses to stimulus location in their receptive field centers. However, it is possible that neuronal representation delays based on population coding may lead to different conclusions. Hence we proceeded to check if we could reproduce the main results of the study presented in Fig. 7 and Fig. 9 using probabilistic population decoding that does not use any response alignment to receptive field centers to compute representation delays. Rather, the moving bar position is decoded based on the population response. Note that this approach was restricted to the results presented in Fig. 7 & Fig. 9 and not used for results in Fig. 10 & Fig.11 because there was an insufficient number of neurons for reliable decoding. We also pooled the two motion directions to obtain a robust estimate of motion latency especially at high speeds where the moving bar traverses the decoded space very quickly giving much fewer trajectory positions to obtain a reliable latency estimate. Similarly, to improve the position decoding under the lower luminance conditions where the neural activity is diminished, we averaged the motion latencies across the two motion directions.

A probabilistic Bayesian decoder (see Methods) was used to estimate the representation delays of the stimuli based on simultaneously recorded single- or multiunit population activity. We assumed that the neurons spike as inhomogeneous Poisson processes that are conditionally independent given the stimulus, and used a decoder trained on flashes to decode moving stimuli. It is well-established that population activity in V1 at a given time is influenced by the location of the bar stimulus and signal conduction and processing delays. This notion is captured in the forward probabilistic model of population activity in Fig. 3. Based on this formalism, a joint distribution of stimulus location, population activity and response delay was obtained (*Eq*.2) from which a posterior probability estimate (*Eq*.5) of a stimulus position can be obtained from the population activity at any given time. Based on the encoding that was learnt from the flash-evoked responses, we decoded the position of the moving bar under different speeds and luminance values. For decoding flashes, we used trials that were not used for encoding to prevent over-fitting. For the luminance modulation experiment, the decoding of bar stimuli of a given luminance was based on encoding obtained from responses to flashes of matching luminance.

The probability of the stimulus position given population activity at different times was computed trial by trial using simultaneously recorded single-units (Fig. 13-A) or multiunits (Fig. 12-A & Fig. 14-A). The resulting position estimates were first averaged across trials and then across sessions (Fig. 12-Fig. 14, B & D). The latency of the peak of the posterior probability (Fig. 12-Fig. 14, C) was taken as the representation delay of the flashes. For the moving bars, first we computed the distance (spatial lag) between the most probable stimulus location and the instantaneous location of the moving bar. Towards this, the trial and session-averaged posterior probabilities (rows in Fig. 12-Fig. 14 D) were aligned (centered) to the instantaneous horizontal positions of the moving bar center (white dots in Fig. 12-Fig. 14 D). For each speed and direction, the aligned probabilities were averaged across the instantaneous positions of the motion trajectory (Fig. 12-Fig. 14, E). The distance between the peak of this aligned probability and the origin gives the spatial lag of the most probable stimulus location. Note that we did not intend to decode the motion speed hence we treated it as a known quantity. The latency of the moving bar representation was then computed by dividing the spatial lag by speed.

**Fig. 12.**
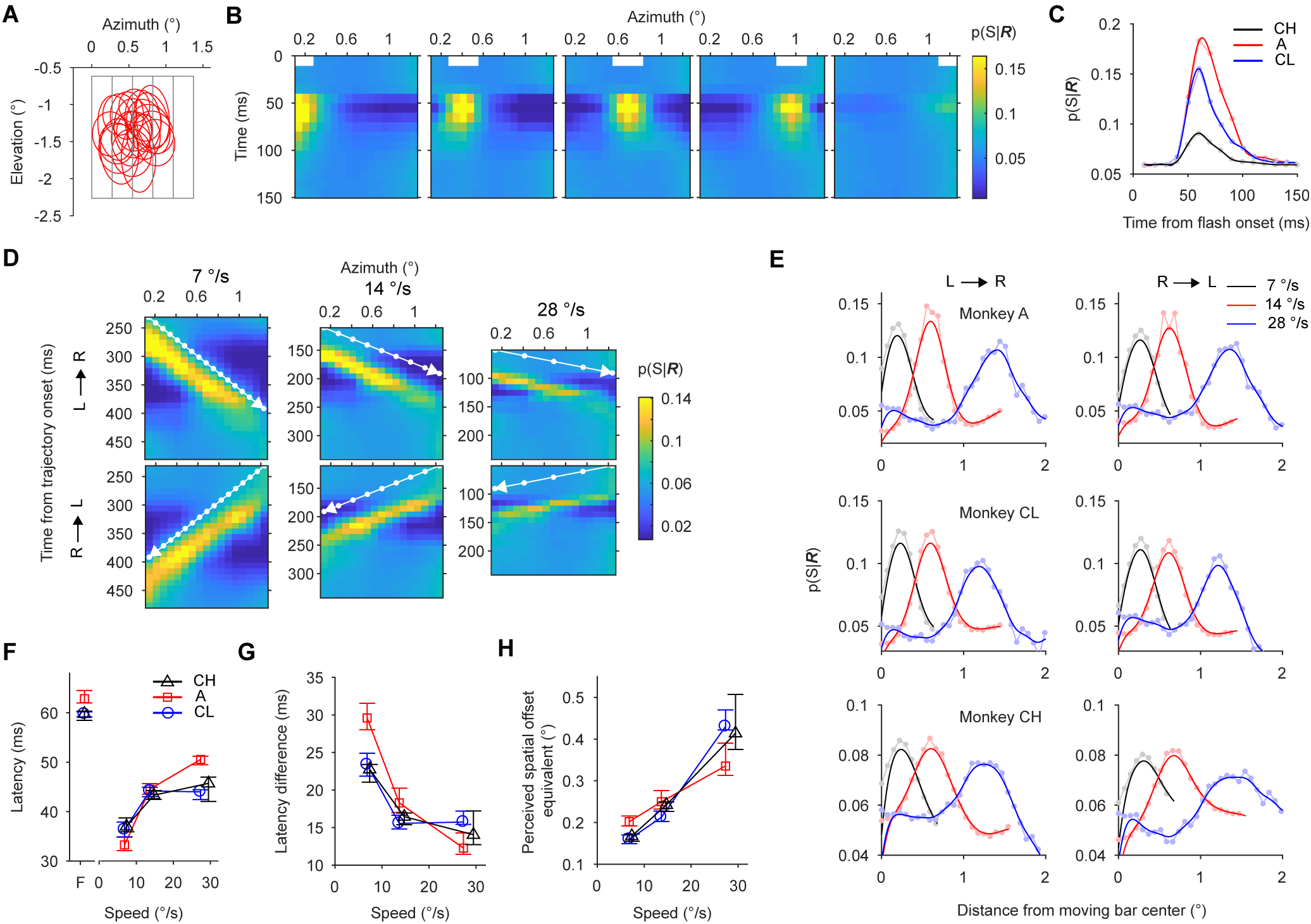
Population multiunit activity decoding of flashes and moving bars and its relationship to flash lag psychophysics under speed manipulation. ***A***, Outlines of receptive fields (red) of simultaneously recorded multiunits from a single representative session. The gray rectangles show the outlines of different flashes presented one at a time. ***B***, Flash decoding results from monkey CL: the white box shown near the top of each panel marks the horizontal position of flash in space and time. Colors of the plot indicate the average (across trials and sessions) probability (*p*(*S*|***R***)) of a horizontal bar position (S) given population activity at a given time (***R***). ***C***, Average probability of flash location, pooled across all flash conditions for individual monkeys (A, CH and CL). ***D***, Average probability of moving bar position for different speeds (panel columns) and directions (panel rows, L→R, motion from left to right) for monkey CL. The white arrows indicate part of the motion trajectory that lies within the flashed region of space. The white dots on the motion trajectory indicate moving bar centers. ***E***, Moving bar probability (rows in panel ***D***) aligned to the instantaneous horizontal position (white dots in panel ***D***) of the moving bar center. For each speed and direction, the aligned probabilities were averaged across the instantaneous positions of the motion trajectory. ***F***, Latency of decoding flash and moving bar locations. Flash (‘F’) latency is the latency of peak of flash location probability in panel ***C***. Moving bar latency for a given direction is the product of the spatial lag of the peaks of moving bar probabilities in panel ***E*** and the inverse of the corresponding speed. Motion latencies were then averaged across directions. Error bars: 95% bootstrap percentile-based plug-in estimate of confidence intervals. ***G***, Speed dependence of the latency difference (Flash minus moving bar latency). Error bars as in ***F. H***, Speed dependence of perceived spatial offset equivalent obtained by the product of the latency difference and speed. Error bars as in ***F***. Colors extremes in B and D are clipped at [0.1, 99.9] percentiles. In ***C*** and ***E***, traces of lighter shades with filled circles correspond to unsmoothed raw data. Median number of trials (sessions) = 289(9), 604(23), 181(10) and median number of multiunits per session (total) = 20(178), 7(166), 19(180) for A, CH and CL respectively.

**Fig. 13.**
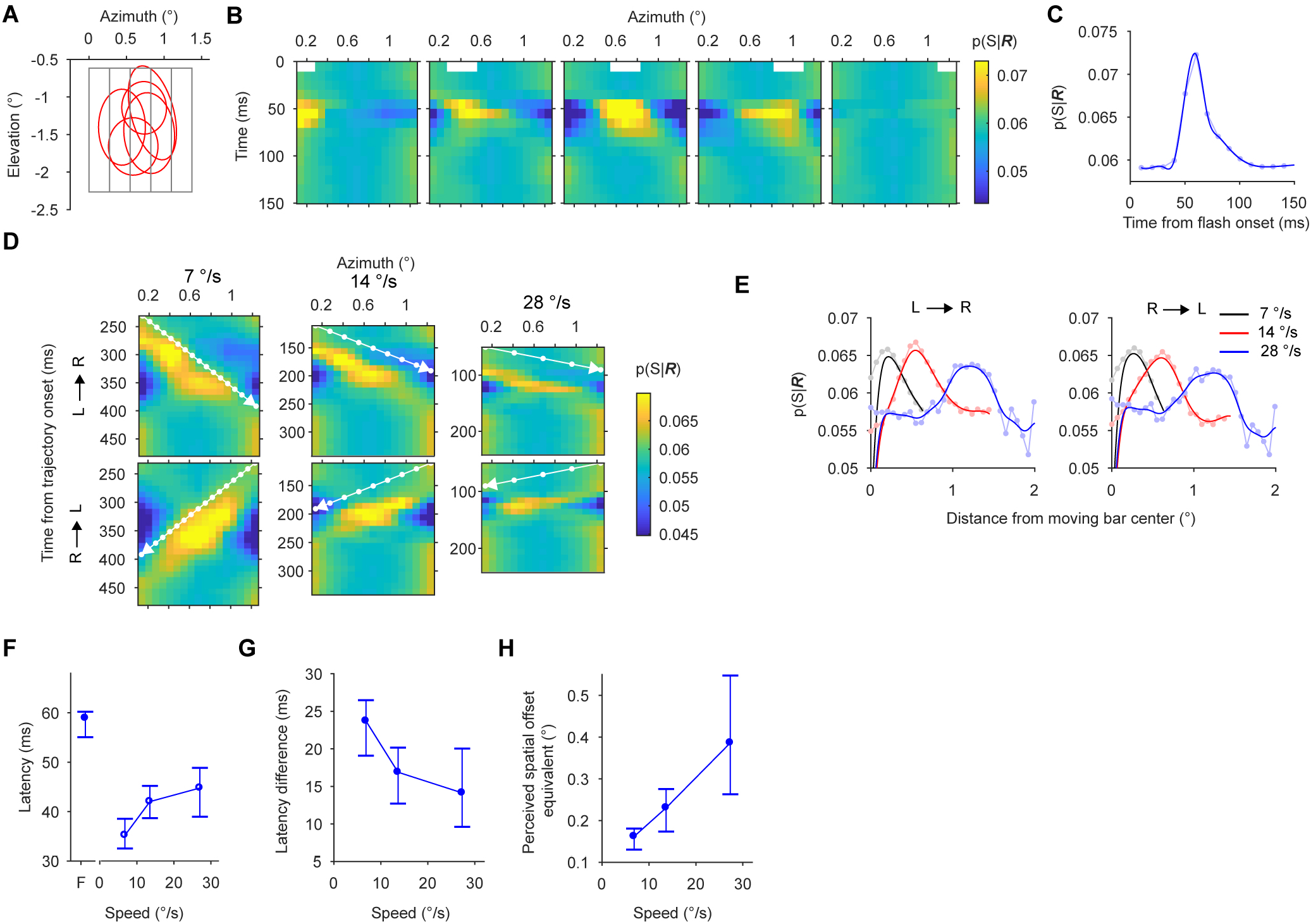
Population single-unit activity decoding of flashes and moving bars and its relationship to flash lag psychophysics under speed manipulation. ***A***, Outlines of receptive fields (red) of simultaneously recorded single-units from a single representative session. The gray rectangles show the outlines of different flashes presented one at a time. ***B***, Flash decoding results from monkey CL: the white box shown near the top of each panel marks the horizontal position of flash in space and time. Colors of the plot indicate the average (across trials and sessions) probability (*p*(*S*|***R***)) of a horizontal bar position (S) given population activity at a given time (***R***). ***C***, Average probability of flash location, pooled across all flash conditions for monkey CL. ***D***, Average probability of moving bar position for different speeds (panel columns) and directions (panel rows, L→R, motion from left to right) for monkey CL. The white arrows indicate part of the motion trajectory that lies within the flashed region of space. The white dots on the motion trajectory indicate moving bar centers. ***E***, Moving bar probability (rows in panel ***D***) aligned to the instantaneous horizontal position (white dots in panel ***D***) of the moving bar center. For each speed and direction, the aligned probabilities were averaged across the instantaneous positions of the motion trajectory. ***F***, Latency of decoding flash and moving bar locations. Flash (‘F’) latency is the latency of peak of flash location probability in panel ***C***. Moving bar latency for a given direction is the product of the spatial lag of the peaks of moving bar probabilities in panel ***E*** and the inverse of the corresponding speed. Motion latencies were then averaged across directions. Error bars: 95% bootstrap percentile-based plug-in estimate of confidence intervals. ***G***, Speed dependence of the latency difference (Flash minus moving bar latency). Error bars as in ***F. H***, Speed dependence of perceived spatial offset equivalent obtained by the product of the latency difference and speed. Error bars as in ***F***. In ***B*** and ***D***, for the set of flash conditions or moving stimuli of a given speed, color boundary was fixed at [0.1, 99.9] percentile. In ***C*** and ***E***, traces of lighter shades with filled circles correspond to unsmoothed raw data. Median number of trials (sessions) = 181(10) and median number of single-units per session (total) = 3(27).

**Fig. 14.**
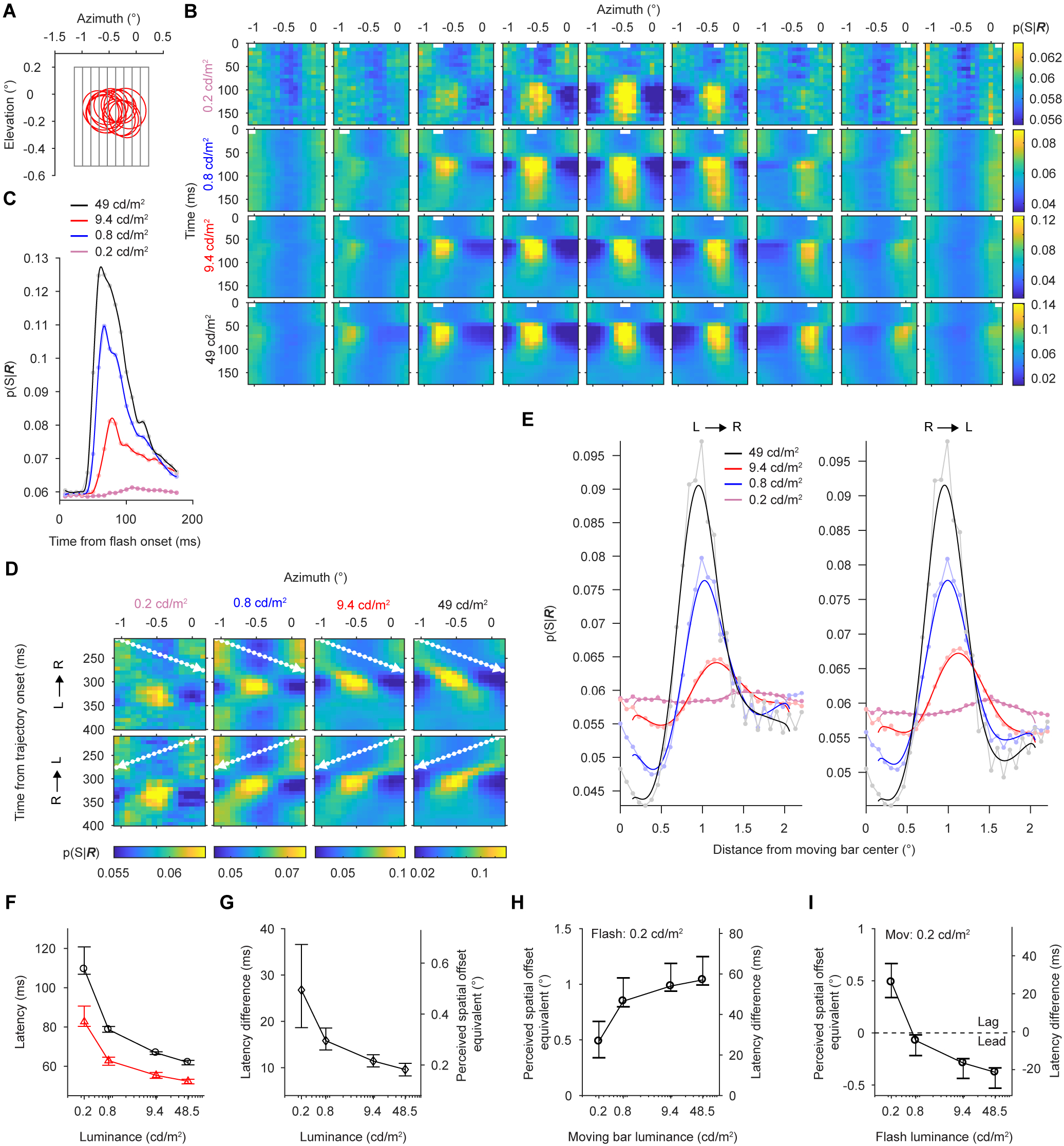
Population multiunit activity decoding of flashes and moving bars and its relationship to flash lag psychophysics under luminance manipulation in monkey L. ***A***, Outlines of subset of receptive fields (red) of simultaneously recorded multiunits from a single representative session. The gray rectangles show the outlines of different flashes presented one at a time. ***B***, Flash decoding results: the white box shown near the top each panel marks the horizontal position of flash in space and time. The luminance of the bars in each row is indicated on the left. Colors of the plot indicate the average (across trials and sessions) probability (*p*(*S*|***R***)) of a horizontal bar position (S) given population activity at a given time (***R***). ***C***, Average probability of flash location, pooled across all flashes of a given luminance. ***D***, Average probability of moving bar position for different luminance values (panel columns) and directions (panel rows, L→R, motion from left to right). The white arrows indicate part of the motion trajectory (speed, 18 °/s) that lies within the flashed region of space. The white dots on the motion trajectory indicate moving bar centers. ***E***, Moving bar probability (rows in panel ***D***) aligned to the instantaneous horizontal position (white dots in panel ***D***) of the moving bar center. For each speed and direction, the aligned probabilities were averaged across the instantaneous bar positions of the motion trajectory. ***F***, Latency of decoding flash and moving bar locations as a function of luminance. Flash (black trace) latency is the latency of peak of flash location probability in panel ***C***. Moving bar latency (red trace) is the product of the spatial lag of the peaks of moving bar probabilities in panel ***E*** and the inverse of the speed. Error bars: 95% bootstrap percentile-based plug-in estimate of confidence intervals. ***G***, Luminance dependence of the latency difference (flash minus moving bar latency, left vertical axis) and perceived spatial offset equivalent (right vertical axis) obtained by the product of the latency difference and speed. ***H*** and ***I***, Latency difference and perceived spatial offset equivalent as a function of moving bar luminance (***F***) for a constant flash luminance (0.2 cd/m^2^) or as a function of flash luminance (***G***) for a constant moving bar luminance (0.2 cd/m^2^). The dotted line in ***G*** separates the luminance conditions that gave rise to perceived spatial offset equivalent corresponding to psychophysically measured flash-lag (‘Lag’) and flash-lead (‘Lead’) conditions. Error bars as in ***B***. In ***B*** and ***D***, for stimulus conditions under each luminance, color bounds were fixed at [0.1, 99.9] percentile. In ***C*** and ***E***, traces of lighter shades with filled circles correspond to unsmoothed raw data. Median number of trials (sessions) = 132(7) and median number of multiunits per session (total) = 39(256).

As reported in Fig. 7B-D, in all three monkeys, based on multiunit population decoding, as speed increased, the motion latency increased (Fig. 12F, p < 0.0005, Bootstrap test), latency difference decreased (Fig. 12G, p < 0.0005, Bootstrap test), and the perceived spatial offset equivalent increased (Fig. 12-H, p < 0.0005, Bootstrap test). From one of the monkeys (CL), we were able to isolate a sufficiently large number of single units, so we were able to verify that the results held true for single well-isolated neurons (Fig. 13 F-H) as well.

For the luminance modulation experiment, we decoded stimulus position for flashes (Fig. 14B-C) and moving bars (Fig. 14D & E) as described above. Again, as found before in Fig. 9, the multiunit population decoding showed that for all luminance values tested, the latency of moving bar was less than that of flashes (Fig. 14 F, p < 0.0005, Bonferroni corrected), latency difference and perceived spatial offset equivalent decreased with luminance (Fig. 14 G, p < 0.0005, Bootstrap test). Similarly, the perceived spatial offset equivalent increased with moving bar luminance when flash luminance was fixed at the lowest value tested (Fig. 14 H, p < 0.0005, Bootstrap test). When the moving bar luminance was fixed at the lowest value tested, the perceived spatial offset equivalent decreased (Fig. 14 I, p < 0.0005, Bootstrap test) changing from being positive (flash-lag) to negative (flash-lead) as the flash luminance level was increased. These results suggest that our conclusions on speed and luminance dependence of latencies and perceived spatial offset equivalents based on individual multiunit responses are consistent with those obtained by population decoding.

## Discussion

Our results show that moving stimuli are processed faster than flashed stimuli in awake macaque V1. In particular, the latency difference between the neural representations of the two stimuli depends on luminance and speed in a way that resembles the perceptual effects of these manipulations in both monkeys (Subramaniyan et al. 2013) and humans (Krekelberg and Lappe 1999; Murakami 2001; Nijhawan 1994; Öğmen et al. 2004; Patel 2000; Purushothaman et al. 1998; Subramaniyan et al. 2013).

Both pre-cortical and cortical mechanisms likely contribute to the observed faster motion processing. These mechanisms potentially include motion induced dynamic shift in the receptive field location and faster conduction/processing of motion signals. Our data cannot distinguish between these two possibilities since both will give rise to a shift in motion response relative to flash response. Motion-induced receptive field shifts have not been reported in the pre-cortical stages in macaques. If found, it would suggest that the labeled line code is not static but more dynamic and will depend on properties of the stimuli. However, there is some evidence for shorter latency of motion signals in the pre-cortical stage - the lateral geniculate nucleus (LGN). In anesthetized cats, it was found that in the different types of LGN cells, the response peak latency for moving bar was shorter compared to that of flashed bar (Orban et al. 1985). Future studies are needed to confirm these findings in monkeys in order to locate the mechanisms underlying the flash lag effect. Cortical processing such as gain control similar to that described in the retina (Berry et al. 1999) and motion-related feedback signals may contribute to dynamic shift in the receptive field location towards the motion direction. For example a recent study (Ni et al. 2014) found that V1 receptive fields in fixating macaques shifted by about 10 % (0.1°) on average in the direction that accounted for the size-distance illusion. Such receptive field shifts if induced by motion can readily explain part of the faster motion processing. Another study that addressed a different illusion called flash-jump illusion also found that V4 neuronal receptive fields shift when the color of one of the bars of an apparent motion sequence changes abruptly (Sundberg et al. 2006). Given that a color change was necessary for such a shift, the implications of their study to the neural mechanisms of flash lag illusion remains unclear.

Faster cortical motion processing could also be achieved by the spreading of subthreshold activity through lateral connections from the currently activated cortical region into the region activated in the future. This spread may facilitate responses by bringing the membrane potential of the target neurons closer to threshold. As a result, those neurons will reach their peak firing earlier, resulting in shorter motion latency. The influence of such subthreshold activity has already been reported in cat V1 in the context of line-motion illusion where the spread of subthreshold activity initiated by one stimulus facilitates the response to a subsequently presented stimulus (Jancke et al. 2004a). Based on this mechanism, it could also be expected that the slower motion would exhibit shorter latency through this mechanism compared to the faster one as there would be more time for the subthreshold activity to spread farther for the slower compared to the faster motion, potentially explaining the speed dependence of motion latency we observed.

We found that the moving bar response peak latency increased with speed. Consistent with our results, conversion of the direction-averaged spatial lag data reported by Jancke et al. (2004) (Fig. 6 in their study) into latency also revealed a similar trend in the speed dependency of motion peak latency. Our data show that latency difference between flash and motion condition decreased with speed. This is in sharp contrast to the constant latency difference that most psychophysical studies assume when interpreting the effect of speed in perceived spatial offset (Krekelberg and Lappe 1999; Murakami 2001; Nijhawan 1994; Whitney et al. 2000). Equivalent latency difference computed from the perceived spatial offsets from a recent psychophysical study (Wojtach et al. 2008) however clearly decreases with speed (Fig. 15) similar to our findings. The discrepancy among the psychophysical studies can be reconciled by noting that Wojtach et al. (2008) used a wide range of speeds (up to 50 °/s) whereas the previous ones used a narrow speed range (up to ~15 °/s), which missed the full trend of the speed effect.

**Fig. 15.**
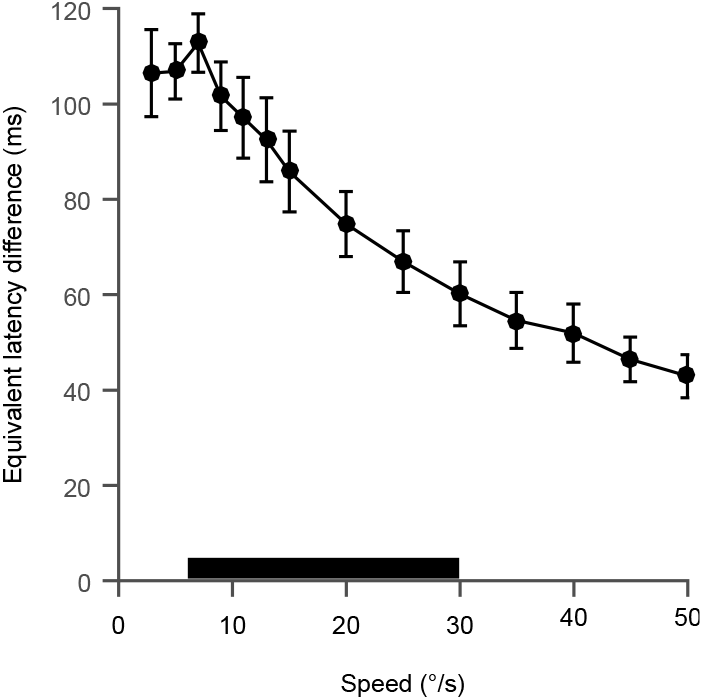
Equivalent latency difference as a function motion speed in humans. The equivalent latency difference (flash temporally lagging the moving object) data points in the plot were computed by dividing the perceived spatial offset by speed reported in Fig. 4 of Wojtach et al. (2008). Error bars are ±1 SEM. The black bar shows the range of speeds used in our physiological experiments.

We found that the perceived spatial offset equivalent depended on speed and luminance (Fig. 7D, Fig. 9C, Fig. 12-Fig. 13, H and Fig. 14G) in line with psychophysical results (Fig. 7E and Fig. 9E). The magnitude of the perceived offset computed from the population decoding method appeared to be closer to the behaviorally measured values than the values computed based on individual multi-unit activity. Interpreting our data conservatively, we think that the perceived spatial offset equivalents we measured in V1 are likely to be smaller than the behaviorally measured values for the following reasons. 1) We measured neural responses from the very first cortical processing stage and the physiological effect may get larger as the information is processed further in the higher cortical areas, 2) the smaller receptive field sizes in V1 may potentially limit the extent to which receptive field shifts can occur in order to reduce motion stimulus representation delays and 3) the monkeys we recorded from did not perform the task and making a relative position judgment may lead to a larger physiological effect. Moreover, we may have also underestimated the discrepancy between the behaviorally measured perceived spatial offset and its neural equivalent because we presented flashes randomly in multiple locations (5-7) for physiology whereas for psychophysics the flash was presented at one (Fig. 9E) or two (Fig. 7E, Fig. 10B-D) fixed locations. Given that predictability of flashes is known to reduce the flash lag effect (Baldo et al. 2002; Brenner and Smeets 2000; Krekelberg et al. 2000; Vreven and Verghese 2005), it is possible that psychophysical measurement of the lag could have been higher if the flashes were equally unpredictable as in our physiology experiments.

It should be noted that the human psychophysical data were collected from two non-naïve subjects whose bias could have an effect on the observed data. We think that this is less likely because our results build upon previously well-established psychophysical results on luminance manipulation (Lappe and Krekelberg 1998; Öğmen et al. 2004; Purushothaman et al. 1998) and are well in accordance with what would be predicted from them. Never the less, further experiments from naïve subjects would be essential to confirm our human psychophysical results.

In our luminance manipulation experiment, we kept the background luminance near zero and changed only the bar luminance. This stimulus configuration, although suitable for mimicking flash lag psychophysical experiment, is not readily comparable to previous physiological studies in V1 that examined luminance or contrast effect on latency using different stimulus configurations (Carandini and Heeger 1994; Gawne et al. 1996; Maunsell and Gibson 1992; Oram 2010; Reich et al. 2001). Despite these stimulus differences, similar to the above studies, we also observed consistent increase in latency when the flash luminance was lowered. The latency for moving bar also increased when the bar luminance was decreased. However, unexpectedly we found that luminance manipulation affected the latency of flash and moving bars differently. The latency profile of the moving bar response was not simply a downward-shifted version of the flash response latency profile. Instead, the increase in latency of moving bar was much less pronounced compared to that of the flash when the luminance was low. With the caveat that the we examined the luminance effect only in a single monkey, these results suggest that moving bars do not suffer as much processing delay as the flashed objects under low luminance conditions and likely invoke different set of mechanisms in bringing out the observed latency effect.

Although several aspects of the flash lag illusion were similar between the monkeys and humans, it was surprising to find that monkeys reported a larger lag for foveofugal motion as opposed to foveopetal motion as found in humans (Kanai et al. 2004; Mateeff et al. 1991; Shi and Nijhawan 2008) and this behavioral effect had a neural correlate in three out of four monkeys. Although species difference could be partly responsible for this, further investigations are needed to fully understand the sources of the discrepancy.

Our data provide two independent lines of evidence supporting the differential latency (DL) model (Öğmen et al. 2004; Patel 2000; Purushothaman et al. 1998; Whitney and Murakami 1998), which predicts shorter time needed for representing moving stimuli. First, as predicted, the perceived spatial offset equivalent computed directly from the latency difference, increased with the speed of the moving bar. Second, the luminance dependence of the flash and motion representation delays (Fig. 9B) is also consistent with the key predictions of DL (Öğmen et al. 2004; Patel 2000; Purushothaman et al. 1998) namely, for a fixed low flash luminance, the perceived spatial offset should increase with moving bar luminance and for a fixed low moving stimulus luminance, progressively increasing the flash luminance should change the flash-lag to flash-lead effect. Our neural data support both predictions (Fig. 9F & G). In addition, latency differences (Fig. 9C) also explained the trend in the luminance modulation of perceived spatial offset using identical luminance for flash and moving bar which we showed in humans (Fig. 9E) for the first time.

According to the motion-biasing model (Eagleman and Sejnowski 2007; Rao et al. 2001), the latencies of flash and moving bar representations are equal, in contrast to what we find in our data. In addition, the illusion arises because “when the brain is triggered to make an instantaneous position judgment, motion signals that stream in over ~80 ms after the triggering event (e.g., a flash) will bias the localization”(Eagleman and Sejnowski 2007). It is unclear exactly in which parts of the brain this ‘biasing’ process is implemented. Also unclear is whether and exactly when V1 spatial representations are altered by this ‘biasing’. Moreover, this model would predict that a neural correlate of the motion-biasing would be observed only when the subjects are asked to make an explicit relative position judgment to decide if the moving and flashed stimuli are misaligned. However, in our main experiments, only a flash or a moving bar was presented in isolation and the animals used in our study were neither trained to make any relative position judgment nor were trained in any other task like the current task; we still found a neural correlate of the illusion in V1. First, these results suggest that reporting relative position judgment is not necessary for observing a neural correlate of the flash lag illusion in visual area V1. Second, they argue against the current version of the motion-biasing model that involves only higher cognitive functions (Eagleman and Sejnowski 2007) and suggest that low level mechanisms underlying the observed latency differences need to be taken into account.

While there is substantial evidence against the spatial extrapolation model at the psychophysical level (Baldo and Klein 1995; Brenner and Smeets 2000; Eagleman and Sejnowski 2000; Lappe and Krekelberg 1998; Purushothaman et al. 1998; Whitney and Murakami 1998), it is possible that spatial extrapolation could be happening at the level of V1. Given that any spatial extrapolation would manifest as a reduction in latency as measured by our method, a full delay compensation as predicted by the model would result in zero response peak latency for the moving bar. However, this was not the case as we found significant delays for the moving bar at all speed and luminance conditions tested. Nevertheless, spatial extrapolation might still hold true in other brain regions or for other sensory systems as shown for auditory motion (Witten et al. 2006).

Irrespective of the model of the flash lag illusion, if the motion representation/perception delays are not ultimately reduced to zero, moving objects will always be mislocalized. Our results suggest that the overall shorter motion latency compared to flashes helps to reduce this mislocalization. Given our results that motion response latencies also change with speed and luminance, how would organisms cope with this in behaviors that require accurate localization of moving objects? One simple and viable solution would be calibration of the sensorimotor integration system. For example, to accurately hit the ball in baseball game, players spend numerous hours in learning (calibrating) to swing the bat at the correct time taking the speed of the ball into account. Hence, the nervous system could in principle learn to respond appropriately to a given moving stimulus condition.

We focused our study on V1 where both flash and motion signals first arrive in the cortex. We showed that moving objects are processed faster in a speed, direction of motion and luminance dependent way compared to suddenly appearing static stimuli. These provide a neural correlate of the flash lag illusion. In this visual area, our data are fully consistent with the predictions of the differential latency model. While the motion-biasing model cannot explain our results, this in itself is not evidence against the model in its entirety. It is possible that the monkeys need to perform the task for the mechanisms proposed by the model to be activated. Visual signals leaving V1 reach a multitude of cortical areas. It is yet to be seen if the differential latency theory would hold in these other areas. Hence further combined behavioral and physiological studies in V1 and subsequent processing stages in the brain are essential to generate additional constraints to narrow down the models.

## Acknowledgements

This work was supported by the National Eye Institute-United States National Institutes of Health (NIH) Grants R01 EY018847, 5-T32-EY07001-37, 5-P30-EY002520-33, the Bernstein Center for Computational Neuroscience (FKZ 01GQ1002), the German Excellency Initiative through the Centre for Integrative Neuroscience Tübingen (EXC307), the Bernstein Award for Computational Neuroscience to PB (BMBF, FKZ 01GQ1601), a Heisenberg Professorship to PB (DFG 5601/4-1) and a German Research Foundation (DFG) grant to ASE (DFG EC 479/1-1). We thank D. Murray, T. Shinn and A. Laudano for technical assistance.

